# Modification of an atypical clathrin-independent AP-2 adaptin complex of *Plasmodium falciparum* reduces susceptibility to artemisinin

**DOI:** 10.1101/621078

**Authors:** Ryan C. Henrici, Rachel L. Edwards, Martin Zoltner, Donelly A. van Schalkwyk, Melissa N. Hart, Franziska Mohring, Robert W. Moon, Stephanie D. Nofal, Avnish Patel, Christian Flueck, David A. Baker, Audrey Odom John, Mark C. Field, Colin J. Sutherland

## Abstract

The efficacy of current antimalarial drugs is threatened by reduced susceptibility of *Plasmodium falciparum* to artemisinin. In the Mekong region this is associated with mutations in the kelch propeller-encoding domain of *pfkelch13*, but variants of other parasite proteins are also thought to modulate the response to drug. Evidence from human and rodent studies suggests that the μ-subunit of the AP-2 adaptin trafficking complex is one such protein of interest. We generated transgenic *Plasmodium falciparum* parasites encoding the I592T variant of *pfap2μ*, orthologous to the I568T mutation associated with *in vivo* artemisinin resistance in *P. chabaudi*. When exposed to a four-hour pulse of dihydroartemisin in the ring-stage survival assay, two *P. falciparum* clones expressing AP-2μ I592T displayed significant and reproducible survival of 8.0% and 10.3%, respectively, compared to <2% for the 3D7 parental line (P = 0.0011 for each clone). In immunoprecipitation and localisation studies of HA-tagged AP-2μ, we identified interacting partners including AP-2α, AP-1/2β, AP-2σ and a kelch-domain protein encoded on chromosome 10 of *P. falciparum*, K10. Conditional knockout indicates that the AP-2 trafficking complex in *P. falciparum* is essential for the fidelity of merozoite biogenesis and membrane organisation in the mature schizont. We also show that while other heterotetrameric AP-complexes and secretory factors interact with clathrin, AP-2 complex subunits do not. Thus, the AP-2 complex may be diverted from a clathrin-dependent endocytic role seen in most eukaryotes into a *Plasmodium-*specific function. These findings represent striking divergences from eukaryotic dogma and support a role for intracellular traffic in determining artemisinin sensitivity *in vitro*, confirming the existence of multiple functional routes to reduced ring-stage artemisinin susceptibility. Therefore, the utility of *pfkelch13* variants as resistance markers is unlikely to be universal, and phenotypic surveillance of parasite susceptibility *in vivo* may be needed to identify threats to our current combination therapies.

## INTRODUCTION

Antimalarial drugs are indispensable components of the global strategy for malaria control and elimination ^1^. Clinical treatment failure with artemisinins, drugs characterised by very short serum half-lives in the treated host, has been demonstrated throughout the Greater Mekong subregion ^2–7^, and there is some evidence of decreasing effectiveness of artemisinin combination therapies (ACT) in Africa ^8–12^. Delayed parasite clearance in Southeast Asia has been linked to several mutations in the propeller-encoding domain of the *pfkelch13* (K13) gene ^6, 13^. However, these mutations have not been observed in parasite populations circulating in Africa, and the function of K13 and its role in resistance remains unclear ^9,11^. Other *Plasmodium* loci have been shown to contribute to artemisinin susceptibility in laboratory studies, including the *P. falciparum* homologue of the actin-binding protein Coronin ^14^. Given that variants of at least two different proteins modulate parasite survival under artemisinin pressure, understanding of any genetic and cellular factors that affect artemisinin susceptibility is desirable for preservation of ACT efficacy worldwide.

Among other candidate loci, the medium (µ) subunit of the AP-2 adaptin trafficking complex (AP-2µ) was first associated with artemisinin susceptibility by Hunt *et al* in multi-drug resistant lineages of *P. chabaudi*, derived by experimental drug selection in rodent hosts ^15^. Linkage analysis identified a mutation encoding PcAP-2µ(I568T) in an artemisinin resistant parasite population when compared with sensitive progenitors ^16^. While subsequent studies failed to identify an orthologous *P. falciparum* AP-2µ(I592T) mutation in human infections, evidence was found for directional selection for a PfAP-2µ(S160N) variant in ACT-treated patients^9^. *In vitro* over-expression suggested this naturally-occurring PfAP-2µ mutation may impact some drug responses, but its role in artemisinin susceptibility remains unclear ^17^. Given the adaptation of the CRISPR-Cas9 system to *P. falciparum* and new data implicating intracellular traffic in the complex artemisinin resistance phenotype, we set out to clarify the impacts of these mutations on *in vitro* artemisinin susceptibility and to probe the underlying biology of the AP-2µ protein in the parasite cell.

In virtually all other eukaryotes, adaptin complexes consist of a heterotetramer and mediate endocytosis and post-Golgi inter-compartment vesicular traffic of receptor-bound cargo ^18^. Each adaptin complex consists of four subunits: two large (α/γ/δ/ε and β), one medium (µ), and one small (σ) ^19^. In particular, the µ subunit interacts with cargo molecules and vesicular membranes and has important regulatory phosphorylation sites ^19,20^. The AP-2 complex is recognised as participating in clathrin-mediated endocytosis from the plasma membrane to the early endosomal compartment in most eukaryotes ^20, 21^. To date there have been no experimental studies of adaptins in *P. falciparum* or other Apicomplexa, and AP-2 is predicted to be involved in endocytosis of haemoglobin from the host cell in *P. falciparum* ^16, 22,23^.

Although the mechanism of artemisinin action in the parasite remains elusive, haemoglobin metabolism is believed to activate the endoperoxide bridge in the artemisinin molecule, required for the drug’s cytotoxic effect ^24-26^. This process may thus link drug activation to trafficking of haemoglobin into the parasite from the host cell cytoplasm. *In vitro* artemisinin resistance is currently defined by survival and expansion of parasite cultures after brief drug exposure during the ring stage, mimicking artemisinin pharmacokinetics *in vivo* ^27^. Reports suggest that activated artemisinin induces disseminated oxidative stress and that resistant parasites have enhanced responses to counter the impact of protein damage and redox stresses centred around the endoplasmic reticulum ^28, 29^. How resistance-associated factors, including variant K13, Coronin, or AP-2μ, contribute to these ring-stage-specific mechanisms is still under investigation.

Here we have investigated the effect of *pfap2mu* mutations on artemisinin susceptibility by generating parasite lineages harbouring I592T and S160N PfAP-2µ mutations via Cas9 genome editing. We then explore the biology of the wildtype PfAP-2µ protein, uncovering unexpected divergences from canonical eukaryotic trafficking biology.

## RESULTS

### A variant of AP-2µ mediates *P. falciparum* ring-stage survival during artemisinin pressure *in vitro*

Two single-nucleotide polymorphisms, encoding PcAP-2µ(I568T), identified in *P. chabaudi*, and PfAP-2µ(S160N), identified in human *P. falciparum* infections, have been suggested to modulate sensitivity to artemisinin in *Plasmodium* ^9, 16^. Using Cas9 editing, we generated clonal *P. falciparum* lines endogenously expressing the orthologous I592T and the naturally-occurring S160N PfAP-2µ mutations on the chloroquine-sensitive 3D7 background (Fig. 1A-B). Transgenic parasites with the integrated mutations were obtained within three weeks of transfection and cloned (Fig. 1B).

**Figure 1.**
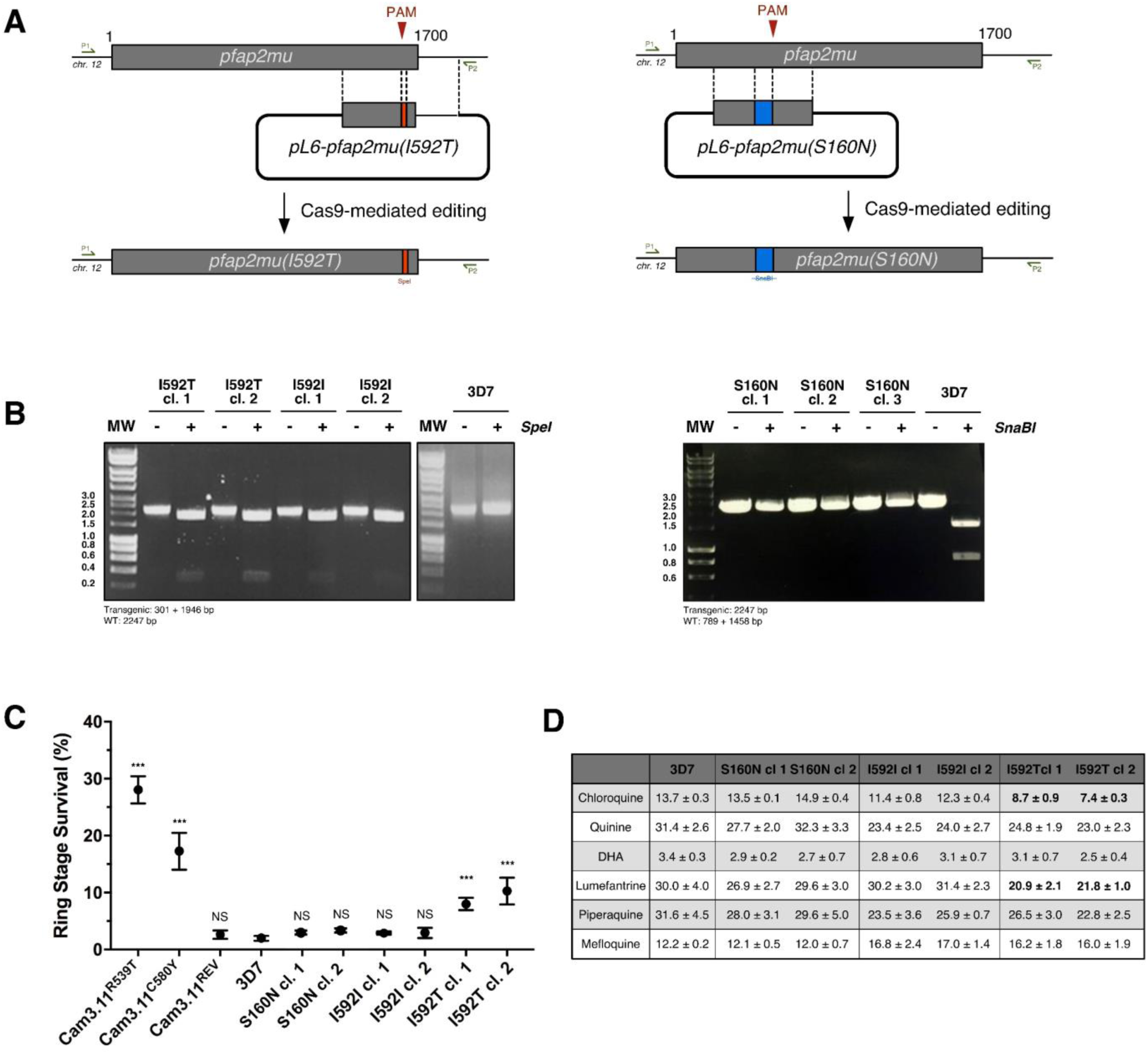
Gene-editing of *pfap2mu* and artemisinin susceptibility by RSA^4h^. **A** CRISPR-Cas9 editing was used to install the *pfap2mu(I592T)* variant codon (red, left) and *pfap2mu(S160N)* variant codon (blue, right) into the endogenous locus. The pictured homologous repair constructs were modified to also introduce the I592I wild-type codon in the context of silent mutations ablating the Cas9 PAM site, as a control for their impact on phenotype. Primers used for genotype mapping anneal outside of the homologous repair template as depicted (P1 and P2; Suppl. Table 1). Recodonised sequence near alternate I592 codons (red) also includes a *Spe*I restriction site, and recodonised sequence near S160 codons (blue) ablates an *SnaBI* restriction site for downstream PCR-RFLP mapping. **B** Clones of parasite lines expressing AP-2µ(I592T), AP-2µ(I592I), and AP-2µ(S160N) were generated by limiting dilution and confirmed by PCR-RFLP genotype mapping (with *Spe*I or *SnaBI* restriction endonuclease) and Sanger sequencing. Amplification of the locus with P1 and P2 produces a 2247 bp fragment; For I592 transgenic parasites, *Spe*I digestion of an amplicon containing the transgenic locus liberates a 301 bp fragment. The wild-type amplicon is not cleaved. For S160 transgenic parasites, *SnaBI* does not cleave the transgenic amplicon, while wild-type amplicons liberate a 789 bp fragment. MW: fragment length in kilobases. **C** RSA^4h^ susceptibility to artemisinin (DHA) of mutant and parental parasite lines, presented as percent survival, compared to untreated control, following a 4h pulse of 700 nM dihydroartemisinin. The Cam3.II-family of parasite lines harbour *pfkelch13* mutations (as indicated)^13^. Cam3.II^REV^ expresses wild-type *pfkelch13*. Mean of at least four biological replicates is shown for each line, each performed in technical duplicate, with standard error. Each technical replicate enumerates 100,000 gate-stopping events. P values derived from Mann-Whitney non-parametric test, comparing each parasite line to 3D7. **D** EC_50_ estimates (nM) for clones of each mutant and the 3D7 parent were derived from 48h exposures to the drugs named across a range of concentrations. Each data point shows the mean of at least four biological replicates, performed in technical duplicate, with standard error. Bold face indicates P < 0.05 (Mann-Whitney U test), compared to EC_50_ estimate for 3D7. NS: not significant; ***: P < 0.005.

Currently, the only validated *in vitro* measure of artemisinin susceptibility is the ring stage survival assay (RSA) ^27, 30-32^. Thus, we examined the survival of ring-stage parasite cultures, two clones of each mutant line, to a 4h pulse of 700 nM DHA using a modified RSA pulse protocol first described and validated by Yang *et al* ^32-33^. Compared to 3D7, the two independent clones harbouring the I592T mutation displayed significantly higher RSA^4h^ survival rates of 10.3% (P = 0.0011) and 8.0% (P = 0.0011), respectively, contributing an approximately 4-5-fold increase in RSA survival when compared to wildtype progenitors. Clones harbouring the S160N mutation displayed equivalent survival after the artemisinin pulse when compared to wildtype progenitors. For comparison, Cambodian clinical isolates harbouring either the circulating R539T or C580Y K13 mutations displayed 28.0% and 17.3% RSA^4h^ survival in our hands, respectively (Fig. 1D), contributing 10.6-and 6.5-fold increases in survival, respectively, compared to an isogenic K13^wt^ Cambodian isolate.

Standard 48hr EC_50_ estimates for quinine, piperaquine, and mefloquine susceptibility were not affected in clones of either *pfap2mu*-variant lineage, but we did observe modest, but statistically significant, potentiation (decreased EC_50_) of lumefantrine and chloroquine in both I592T clones (Fig. 1C). As previously demonstrated for other artemisinin resistant lineages, 48h EC_50_ estimates for artemisinin were unchanged in the I592T clones, compared to wildtype progenitors. We also generated a transgenic parasite line expressing the wildtype I592I codon in the context of silent “shield” mutations required for obliterating the Cas9 PAM site in the *pfap2mu* locus used during I592T editing to confirm the effects observed in the RSA and 48h assays were directly attributable to the I592T SNP. As expected, two clones of this I592I lineage demonstrate equivalent sensitivity in all assays when compared to 3D7. These data demonstrate that AP-2μ mutation can confer significant ring-stage protection from artemisinin in an otherwise drug-susceptible *P. falciparum* strain *in vitro*.

### AP-2μ is localised to a cytoplasmic compartment

A mechanistic understanding of I592T-mediated artemisinin resistance first requires knowledge of the biology of the wildtype PfAP-2µ protein. To generate this information, we sought to characterise the cellular location, essentiality and interactome of this protein. We expected that AP-2µ would be involved in clathrin-mediated endocytosis from the plasma membrane based on eukaryotic trafficking dogma ^19^. Thus, using Cas9 editing, we installed a tandem triple haemagglutinin (3xHA) epitope tag onto the C-terminus of the protein (Fig. 2A-C) and examined the localisation of AP-2µ-3xHA by IFA across the asexual lifecycle in two clones (AP-2µ-3xHA_c1 and AP-2µ-3xHA_c2). AP-2µ-3xHA distribution was similar in both clones, so only the data from AP-2µ-3xHA_c1 are presented. Contrary to our expectations that AP-2µ-3xHA would decorate the parasite plasma membrane, consistent with a role in endocytosis, we observed that, throughout the asexual cycle, AP-2µ localises to additional punctate structures in the parasite cytoplasm (Fig 2D). In the ring stage, AP-2µ-3xHA typically appears as a single focus, near the parasite plasma membrane. As the cell develops, foci increase in number and localise to a cytoplasmic compartment in trophozoites (Fig. 2D). During schizogony, the AP-2μ-labelled structure multiplies and segments into each daughter merozoite (Fig. 2D). To probe the observed punctate localisation of AP-2µ further, its cellular distribution was examined with respect to a panel of representative organelle markers. However, the distribution of AP-2µ signal did not correlate well with that of the ER, Golgi complex, or the apicoplast during development or with the apical secretory organelles during schizogony (Suppl. Fig. 1). When compared with brightfield, a focus of AP-2µ was often in close apposition to the digestive vacuole, but these structures were never observed to overlap (Fig. 2D, middle column).

**Figure 2.**
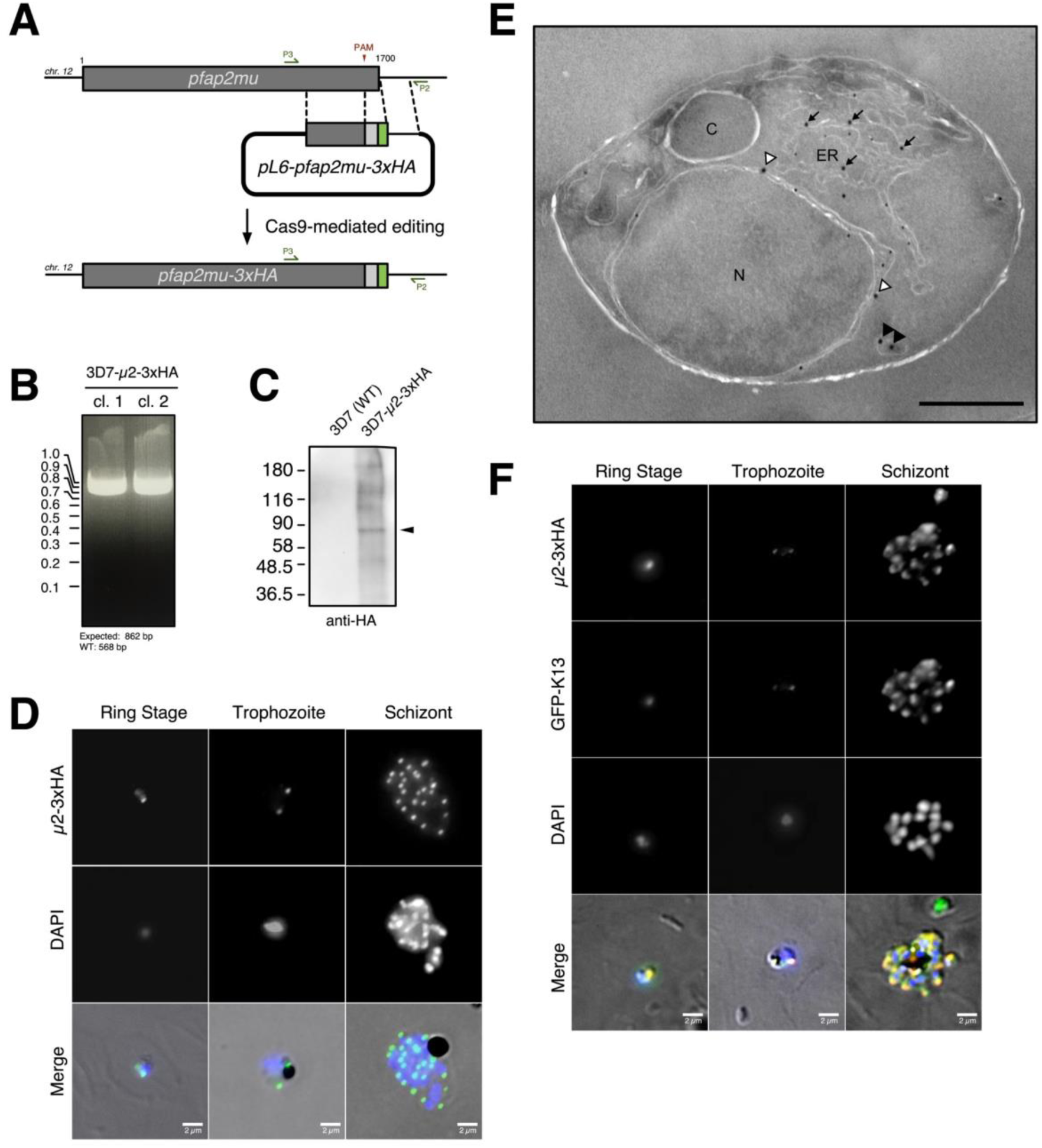
*Plasmodium falciparum* AP-2µ is localised to a non-canonical cytoplasmic compartment. **A** Modifications to the homologous repair construct used to install AP-2µ variants (Fig. 1) to fuse a tandem triple haemagglutinin tag (3xHA) onto the C-terminus of AP-2µ. **B** PCR-based genotyping of two parasite clones harbouring AP-2µ-3xHA in place of the endogenous AP-2µ allele. Amplification of the integrated transgenic *pfap2mu* locus with P3 and P2 (annealing sites annotated) produces an 862 bp fragment. Genotypes were confirmed by Sanger sequencing of the PCR products shown. **C** Anti-HA western blot confirming expression of the desired fusion protein (approx. 78 kDa) in mixed-stage lysates, compared to wild-type parental 3D7. MW presented in kDa. **D** Localisation of AP-2µ-3xHA (green) across the asexual life cycle by anti-HA IFA, counter-stained for parasite DNA with DAPI (blue). Images shown are representative of more than 100 cells examined at each stage; merge is the superimposition of each channel on a brightfield image. **E** Immunoelectron micrograph of a representative young intra-erythrocytic trophozoite. AP-2µ-3xHA parasites probed with an anti-HA rabbit antibody and a secondary antibody 18 nm gold conjugate. Protein disulphide isomerase (PDI), a marker for the parasite ER, is detected by an anti-PDI mouse antibody and a secondary conjugated to 12 nm gold particles. C: cytostome; N: nucleus; ER: endoplasmic reticulum; White Arrowheads: AP-2µ at nuclear membrane; Black Arrowheads: AP-2µ associated with vesicles; Black Arrows: AP-2µ near the ER. Scale bar: 500 nm. **F** Localisation of AP-2µ-3xHA (false-coloured green) with respect to ectopically-expressed GFP-K13 (false-coloured red) across the asexual life cycle by IFA. Representative images of more than 100 observed cells is shown.

To better characterise the localisation of AP-2µ, we performed immunoelectron microscopy (immuno-EM) on thin sections of trophozoites expressing AP-2µ-3xHA. In analysis of 66 micrographs of single parasite-infected erythrocytes, gold particles detecting anti-HA antibodies bound to AP-2µ-3xHA were observed at the nuclear membrane and ER (73.8% of micrographs), in vesicles in the cytosol (37.9%), in close proximity to tubular cytosolic structures (93.6%), near the food vacuole (5.8%), and, rarely, at the parasite plasma membrane (4.2%) (Fig 2E, Suppl. Table 1). Immunostaining against the endoplasmic reticulum (ER)-resident protein disulfide isomerase (PDI) suggests that these tubular structures may be cross-sections of distal extensions of the ER (Fig. 2E). Co-staining against Rab5B, an effector of endosomal transport, suggests at least some of the cytosolic AP-2µ associates with Rab5B^+^ vesicles (Suppl. Fig. 2). Parasites expressing AP-2µ-2xFKBP-GFP showed a similar localisation by immuno-EM (Suppl. Fig. 3, Suppl. Table 1), but GFP fluorescence was too faint to reliably observe in live cells.

Because *Plasmodium* parasites lack a stacked Golgi apparatus, differentiating the ER and Golgi by EM is difficult. Therefore, AP-2µ-3xHA parasites were treated with brefeldin A (BFA), a fungal metabolite and fast-acting inhibitor of ER-to-Golgi secretory traffic. Upon stimulation with BFA, proteins localised to, or trafficked via, the Golgi relocalise to the ER. Previous studies examining intracellular traffic in *Plasmodium* have shown that parasite cultures can be treated with 5 µg/ml BFA for up to 24h while remaining viable ^34, 35^. After treating ring stage parasites with 5 µg/ml BFA for 16h, AP-2µ-3xHA staining significantly co-localised with staining observed for plasmepsin V, a luminal ER protease, suggesting AP-2µ is localised to a secretory membrane. AP-2µ and plasmepsin V staining were distinct in solvent-treated controls. (Suppl. Fig. 4).

In recent studies, K13 has been localised to conspicuous membranous structures in the cytosol ^36^, and these superficially resemble the structures labelled by AP-2µ here (Fig. 2 D, E). Given the apparent similarity in cellular distribution and the importance in ring-stage artemisinin susceptibility *in vitro* of both molecules, we hypothesised that AP-2µ and K13 localise to the same cytosolic compartment. We overexpressed an episomally-encoded N-terminal GFP-K13 fusion protein in our AP-2µ-3xHA-expressing line and observed a striking similarity in signal distribution between the K13 and AP-2µ in fixed ring and schizont stages (Fig. 2F). The GFP-K13 signal resembled what has previously been reported ^36^.

### PfAP-2µ is required for asexual replication

Given these results demonstrating an unexpected cytoplasmic localisation for AP-2µ, we sought to better characterise the protein’s functional importance. Previously results suggested that AP-2µ is required for asexual replication, but its role in asexual development is unclear. Therefore, we deployed the inducible DiCre system described by Collins *et al* to study the phenotypic effects of conditional *pfap2mu* knockout (KO) *in vitro* ^37^. Specifically, the Cas9 donor constructs described above were modified to install both a *loxP* site into the 3’ UTR immediately after the *pfap2mu* stop codon and the *loxP-*containing *pfsera2* intron described by Jones *et al* into the 5’ end of the gene, 261 bp into the CDS, such that Cre-mediated excision removes the majority of the coding sequence, including the 3xHA tag. These constructs were transfected into 3D7 parasites constitutively expressing dimerisable Cre recombinase ^38^ (Fig. 3A).

**Figure 3.**
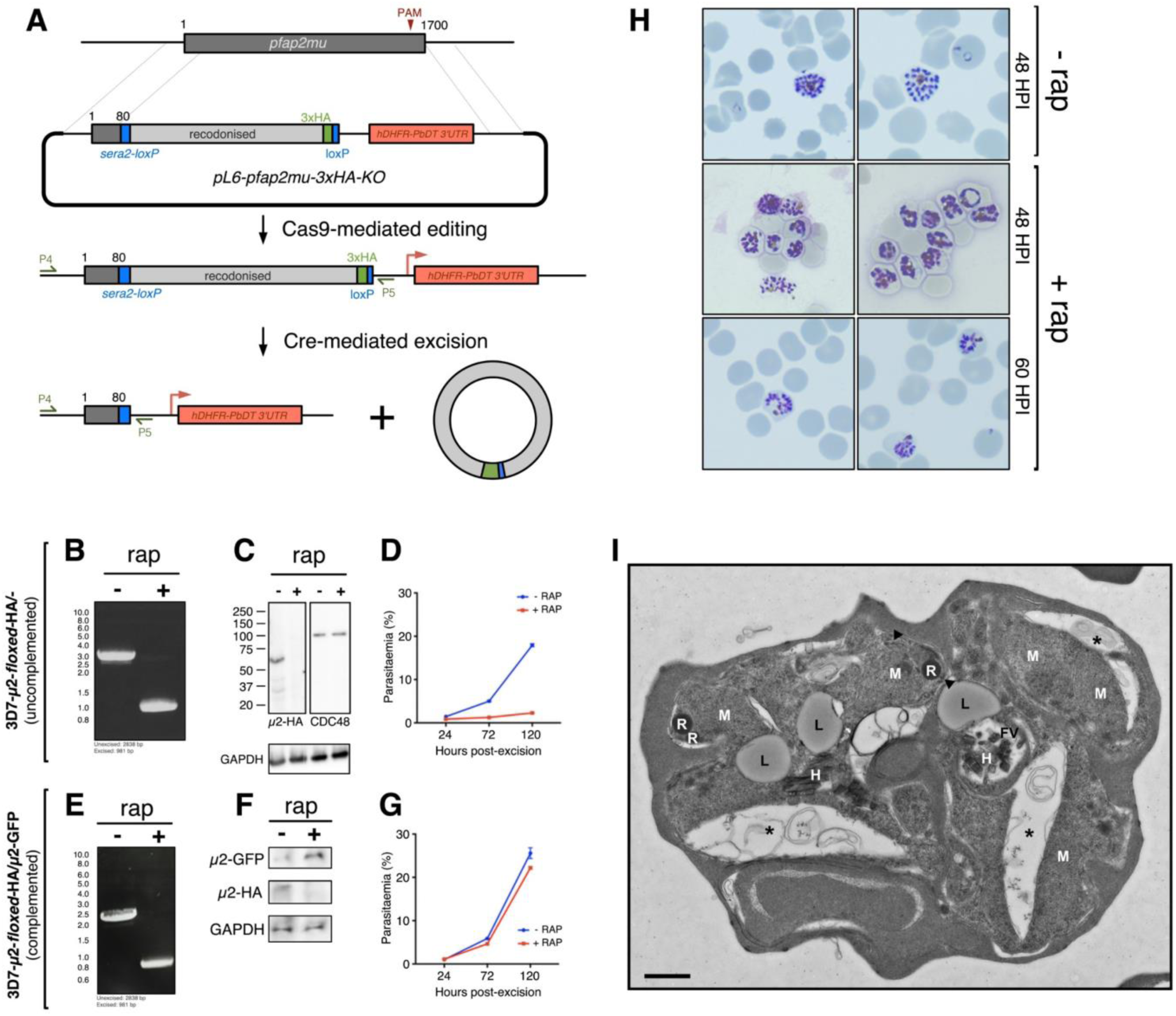
AP-2µ is required for asexual replication and schizont maturation. **A** Schematic for the integration of *loxP* recombination elements into the endogenous *pfap2mu* locus of a parasite line constitutively expressing a split-Cre recombinase^33^. Addition of rap initiates Cre dimerisation and excision of the *loxP*-flanked (*floxed*) region of *pfap2mu* on chromosome 12. **B** PCR confirmation of rap-induced excision of *floxed* region by PCR using primers P4 and P5 (panel A). **C** Western blot confirmation that excision of *floxed pfap2mu* causes a loss of AP-2µ-3xHA protein (within 24h) but has no effect on levels of CDC48 protein. MW presented in kDa. **D** Parasite multiplication in the 3D7-AP-2µ-*floxed*-3xHA line across 2.5 cell cycles, with and without rap induction of Cre. Mean parasitaemia (normalised to 0.25% starting parasitaemia) with standard error is shown at each time point. Each data point represents the average of at least three biological replicates. **E** PCR confirmation of rap-mediated *pfap2mu* excision in 3D7-AP-2µ-*floxed*-3xHA parasites transfected with an episome encoding *cam-*AP-2µ-GFP. Construction of this complementation plasmid is described elsewhere (R. Henrici, PhD Thesis, University of London 2018). **F** Western blot confirmation that excision of chromosomal *pfap2mu* from 3D7-AP-2µ-*floxed*-3xHA/*cam-*AP-2µ-GFP parasites causes a loss of AP-2µ-3xHA protein but does not prevent episomal expression of AP-2µ-GFP. **G** Parasite multiplication in the 3D7-AP-2µ-*floxed*-3xHA/*cam-*AP-2µ-GFP line across 2.5 cell cycles, with and without rap induction of Cre. Mean and standard error are shown as in panel D. **H** Giemsa staining of 3D7-AP-2µ-*floxed*-3xHA schizonts, with and without rap treatment. **I** Electron micrograph of 3D7-AP-2µ-*floxed*-3xHA schizonts, with and without 1h ring-stage treatment with 10 nM rap. Micronemes at the apical end of developing merozoites are labelled with arrowheads; asterisks indicate membrane separation. FV: food (digestive) vacuole; H: haemozoin; L: lipid body; M: merozoite; R: rhoptry. Scale bar: 500 nm.

**Figure 4.**
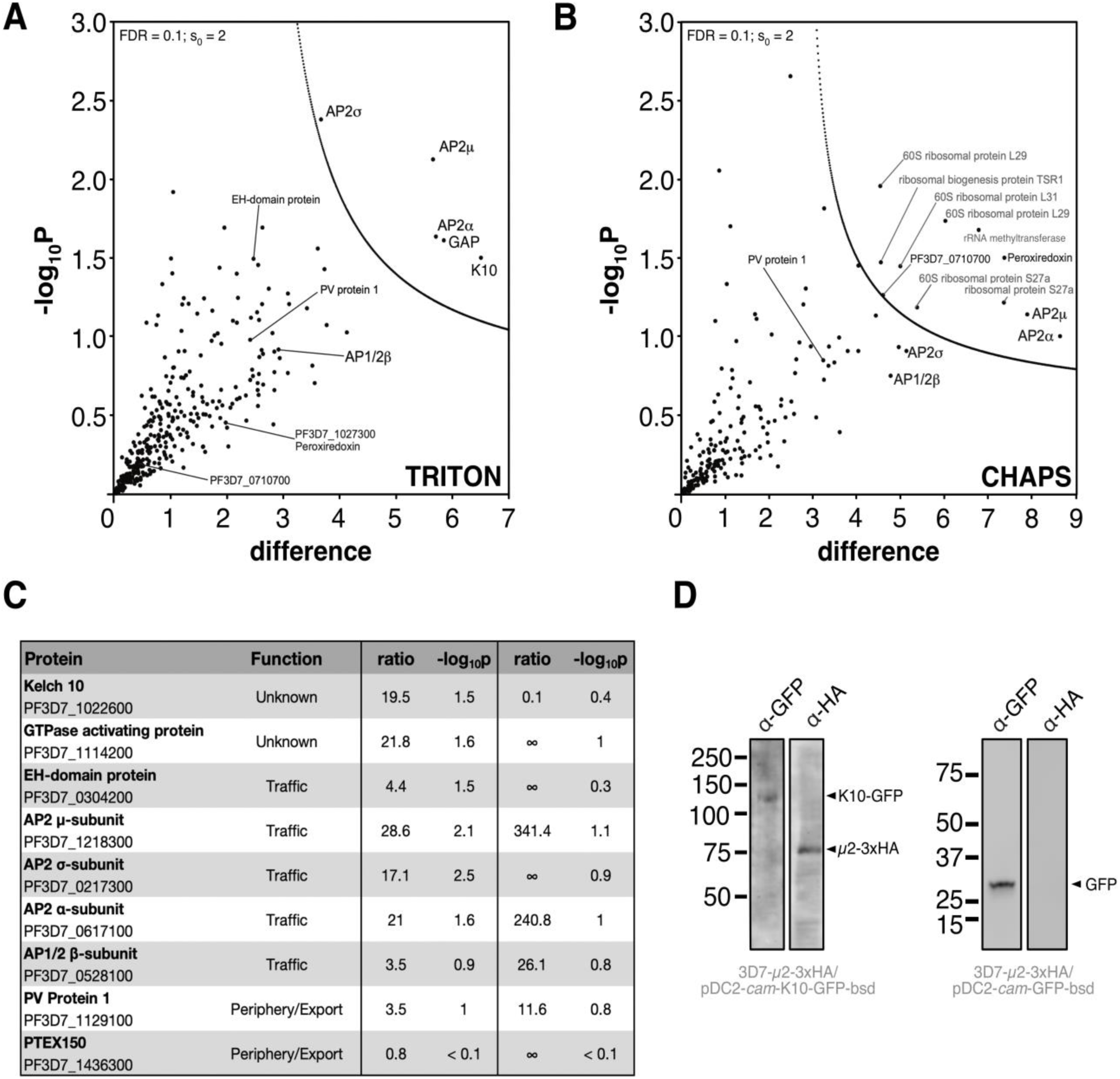
Identification of AP-2µ *-*interacting proteins. **A** Volcano plot from *p*-values versus the corresponding *t*-test difference of proteins identified by immunoprecipitation (IP) in Triton buffer. Cut-off curves for statistically significant interactors (dotted curve) were calculated from the estimated false discovery rate (for details see Methods). Selected hits are labelled (potentially non-specific interactors are in grey). **B** Volcano plot for proteins identified by IP in CHAPS buffer. **C** Table of selected identified interactors listing functional annotation, enrichment ratios (compared to controls; see Materials and Methods) and negative log_10_ of corresponding *p*-values for Triton & CHAPS buffers, respectively. Additional hits are listed in Suppl. Table 2. **D** *pfk10-*GFP (left panel) or GFP alone (right panel), driven by the calmodulin promoter, were expressed episomally in 3D7-AP-2µ-3xHA parasites and immunoprecipitated with α-GFP antibody coated magnetic beads. Western blots of fractionated proteins are shown, probed with either α-GFP or α-HA antibodies. MW presented in kDa.

Ring stage cultures of the 3D7-AP-2µ-*floxed*-3xHA parasites were treated with 10 nM rapamycin (rap) in DMSO for 30 min to dimerise the split Cre recombinase and trigger *pfap2mu* excision. Genomic DNA was extracted 24h after this treatment, and PCR confirmed near-complete excision of the *floxed* region of *pfap2mu* (Fig. 3B), resulting in ablation of AP-2µ protein expression (Fig. 3C). Daily parasite counts by FACS revealed that induced-KO of *pfap2mu* prevented parasite replication within a single asexual cycle, without appreciable recovery over multiple cycles (Fig. 3D), showing that *pfap2mu* is required for asexual replication *in vitro*. Importantly, the Cre-mediated endogenous *pfap2mu* KO was fully complemented with an episomally-expressed copy of AP-2µ-GFP under a constitutive promoter (Fig. 3E-G; Suppl. Fig. 5). These results also support our localisation studies, indicating that disrupting AP-2µ is lethal, and C-terminal AP-2µ tags are thus tolerated by the parasite.

**Figure 5:**
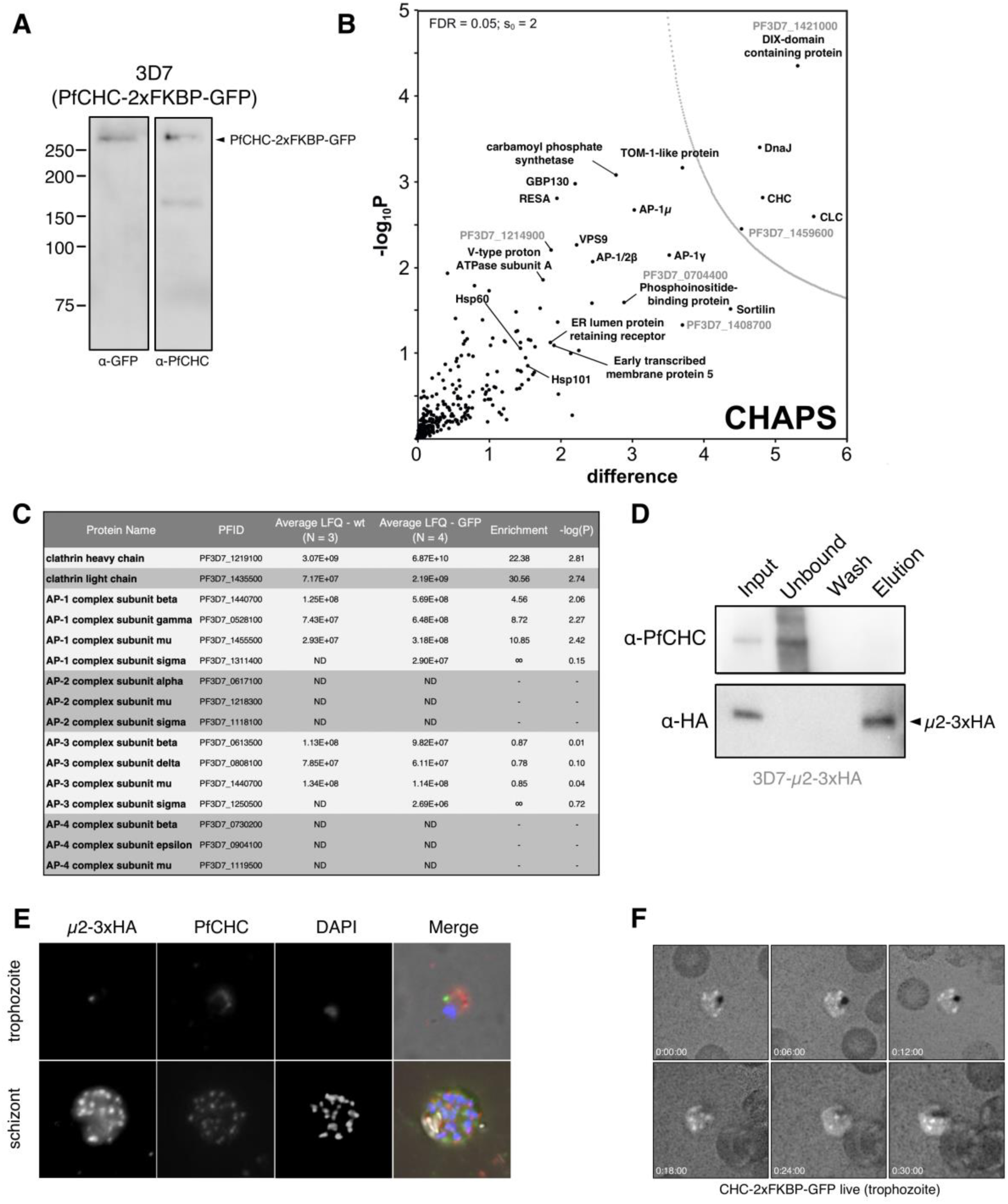
*P. falciparum* AP-2µ does not interact with clathrin heavy chain. **A** Western blot of mixed stage 3D7-CHC-2xFKBP-GFP lysates probed with antibodies; either α-GFP (left) or α-PfCHC (right). **B** Volcano plot from *p*-values versus the corresponding *t*-test difference of proteins identified by α-GFP (nanobody) immunoprecipitation in CHAPS buffer. Cut-off curves for statistically significant interactors (dotted curve) were calculated from the estimated false discovery rate. **C** Table of abundances and enrichment ratio for subunits of all adaptor protein complex subunits identified in the α-CHC-GFP pulldown. ND indicates no peptides were detected that correspond to the listed protein. **D** Western blot of α-PfCHC immunoprecipitation performed on cryomilled 3D7-AP-2µ-3xHA trophozoite lysates. Membrane was probed with α-PfCHC and α-HA antibodies. **E** Maximum intensity projection IFA of representative trophozoite and schizont stages of 3D7-AP-2µ-3xHA parasites, from among at least 100 cells examined at each stage, probed with both α-PfCHC and α-HA antibodies, which are red and green in the merge images, respectively. **F** Representative maximum intensity projection images of time-lapse live microscopic observation of CHC-2xFKBP-GFP in a trophozoite. Each frame represents the passage of 6 minutes.

When examined by Giemsa staining, parasites lacking *pfap2mu* arrest as malformed schizonts (Fig. 3H). These defective schizonts occupy approximately half of the red cell cytoplasm and contain poorly segmented merozoites compared to wild-type schizonts. Despite the reduced size of schizonts in *pfap2mu* knockouts, their DNA content is similar to that of fully segmented, egress-blocked schizonts by FACS (Suppl. Fig. 6). Rarely, rap-treated parasites were observed to undergo egress and invasion despite the addition of rapamycin and probably represent incomplete excision of *pfap2mu* across the cultured population. Supporting this, in our PCR amplification of genomic DNA extracted from these rap-treated cultures, a faint 3kb band, indicative of unexcised full-length *pfap2mu*, was detected (Fig. 3B). This phenomenon of incomplete Cre-mediated excision has been reported previously ^37^.

**Figure 6:**
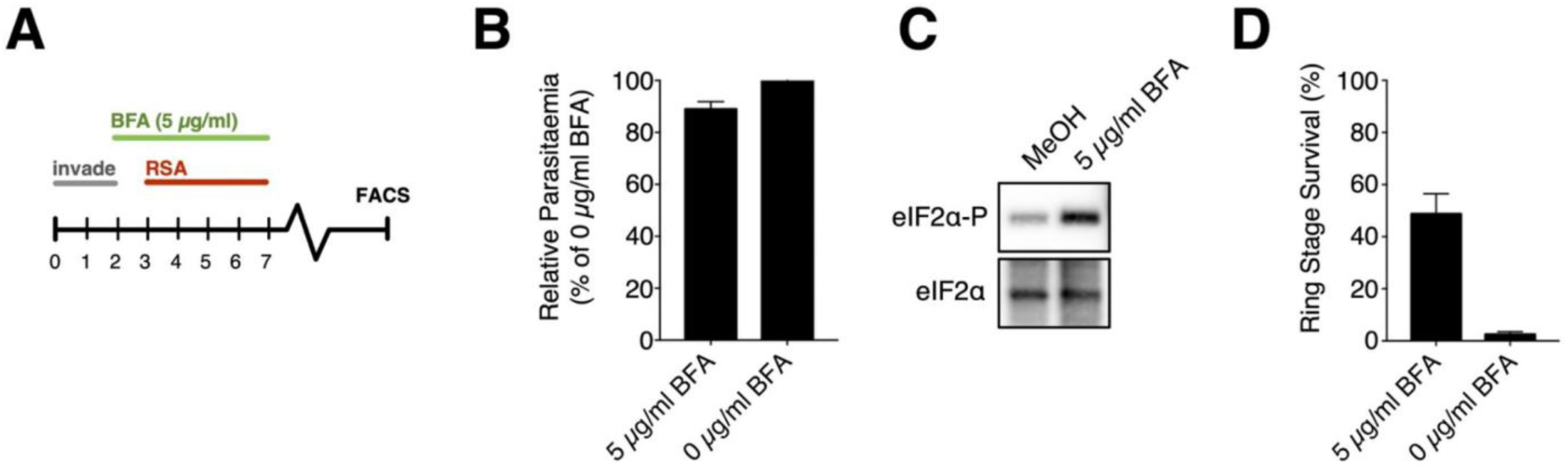
Activation of the cell stress response with BFA induces artemisinin resistance *in vitro*. **A** Schematic representation of BFA^5h^-RSA^4h^ assay. Synchronous cultures of 3D7 parasites received 1h pre-treatment with 5 μg ml^−1^ BFA (green) before addition of DHA for the RSA^4h^ (red). **B** Relative proliferation of wild-type 3D7 parasites treated with 5 μg ml^−1^ BFA^5h^ compared to untreated parasites, in the absence of DHA. **C** Western blot of parasite lysates probed with anti-eIF2α-P or anti-eIF2α, in BFA^5h^-treated compared to solvent-treated cultures. Phosphorylation of eIF2α indicates activation of the cell stress response. **D** Parasite survival (% of untreated) in BFA^5h^-RSA^4h^ and standard RSA^4h^ in 3D7 parasites. Mean survival estimates ± standard error for 3 experiments, each in technical duplicate, is shown.

### PfAP-2µ is required for schizogony

Examining these deformed schizonts by electron microscopy revealed that AP-2µ deletion results in gross morphological defects during schizogony and merozoite biogenesis (Fig. 3I; Suppl. Fig. 6). Merozoites forming within AP-2µ-KO schizonts are highly disorganised and misshapen within the parental schizont plasma membrane, and there are large pockets of schizont cytosol between the malformed merozoites. The schizont plasma membrane seems indistinct, loosely encircling the deformed merozoites with irregular invaginations not seen in wild-type parasites (Fig. 3I; Suppl. Fig. 6). In AP-2µ-KO schizonts, peripheral plasma membranes were consistently poorly preserved during fixation and processing for electron microscopy despite several replicate preparations. These membrane bilayers tended to separate dramatically compared to membranes in wild-type schizonts, and we cautiously attribute this observation to increased membrane fragility in the absence of AP-2μ.

Additionally, electron micrographs revealed a statistically significant accumulation of lipid bodies in the cytosol of AP-2µ KO schizonts (Fig. 3I; Suppl. Fig. 6, black arrows), with some cells having 2, 3 or 4 such bodies (Poisson regression: coefficient 0.597, 95% CI 0.344-0.850; P<0.001; N = 300 treated, 300 untreated). These parasites were also more likely to contain disturbed digestive vacuoles (DV), which often appeared fragmented or elongated (odds ratio 3.18 (95%CI 1.89-5.48; P<0.0001). Consistent with this, haemozoin crystals were occasionally observed in the schizont cytosol. Wild-type and KO trophozoites appear morphologically equivalent (Suppl. Fig. 6), and apicoplasts can be observed in these cells.

Subsequent IFA imaging supported these findings, revealing that *pfap2mu* is required for the genesis of several membrane-bound organelles (Suppl. Fig. 7). Specifically, AMA1, RON4, and CDC48, markers of the micronemes, rhoptries, and apicoplast, respectively, are mis-localised by IFA, despite being detectable at normal levels by western blot (Suppl. Fig. 7, 8). Despite the apparent mislocalisation of AMA1 and RON4 by immunofluorescence (Suppl. Fig. 7), rhoptries and micronemes are visible in some KO schizont EM sections (Fig. 3I), implying a defect in transport rather than organelle biogenesis. Additionally, in KO schizonts, MSP1 antibody staining is disrupted, suggesting that trafficking to the schizont plasma membrane, which invaginates around the nascent merozoites during schizogony, may be disrupted in the absence of AP-2µ. The parasitophorous vacuole, labelled by EXP2, seems to be largely intact though may have unusual discontinuities in some cells. The ER and *cis-*Golgi display no obvious abnormalities in cells lacking AP-2µ (Suppl. Fig. 7). Overall, it is unlikely that these widespread defects are all directly attributable to AP-2µ function. Instead, this complex phenotype may be the result of knock-on effects of AP-2µ deletion affecting downstream effector molecules.

### The AP-2 complex is clathrin-independent and associates with Kelch10

To elucidate some of these possible AP-2µ interacting partners, we immunoprecipitated AP-2µ and performed mass spectrometry. Trophozoite cell lysates of *P. falciparum* AP-2μ-3xHA were prepared using cryomilling and detergent lysis, a cell disruption technique shown to preserve labile protein-protein interactions and generate high resolution interactomes in other organisms, though not previously deployed in *Plasmodium* ^39^. Using both Triton-and CHAPS-containing lysis buffers originally derived for the extraction of clathrin-interacting proteins from *Trypanosoma* species, we lysed frozen trophozoite pellets and performed pulldowns using anti-HA conjugated beads.

As previously discussed, AP-2 is the canonical clathrin-interacting endocytic complex in essentially all other studied eukaryotes, and we expected AP-2µ to interact with clathrin and the other putative AP-2 subunits in *P. falciparum*. Four adaptor complex subunits were enriched under both lysis conditions tested, demonstrating that the AP-2 complex in *P. falciparum* comprises subunits annotated in the genome as AP-2α, AP-2σ, AP-2µ (our tagged bait protein), and AP-1β, previously predicted to be shared between the AP-1 and AP-2 complexes ^40^ (Fig. 4A - C). This latter subunit is therefore designated here as AP-1/2β. Putative nucleotide-dependent regulators of vesicular traffic and a kelch-type propeller domain protein encoded on chromosome 10, K10, were also identified with high confidence under one or both lysis conditions (Fig. 4C; Suppl. Table 3). Though the K10 propeller domain shares little sequence identity with that of K13, *pfk10* has previously been identified as a locus of interest in a genetic study of artemisinin resistance ^41^. K13 was not identified as an AP-2µ-interacting protein in our immunoprecipitation data. Additionally, a putative EH-domain containing protein with homology to EPS15 (a classical AP-2 interacting partner) was enriched in the immunoprecipitation. Strikingly, neither the clathrin heavy chain (CHC) (gene ID: PF3D7_1219100) nor the light chain (CLC) (gene ID: PF3D7_1435500) were enriched in either extraction condition with our HA-tagged AP-2µ, contrary to expectation.

To validate our interactome, we confirmed that AP-2µ-3xHA copurifies with episomally-expressed and immunoprecipitated K10-GFP but not similarly expressed cytosolic GFP (Fig. 4D). We sought to confirm the absence of interacting clathrin by direct IP of CHC from *P. falciparum* AP-2μ-3xHA lysates and western blotting with anti-HA antibodies. We were still unable to find evidence that AP-2µ-3xHA co-purifies with immunoprecipitated CHC (Fig. 5D). Using the same clathrin antibody, first validated on a parasite line expressing PfCHC-2xFKBP-GFP (Fig. 5A), we also found that AP-2µ-3xHA and CHC are localised to separate compartments by IFA (Fig. 5E).

To test robustly the observation that AP-2 does not interact with CHC, we performed a similar IP-MS analysis on a trophozoite preparation of parasites expressing CHC-2xFKBP-GFP. As expected ^42^, all AP-1 complex components were enriched in these PfCHC pulldowns, including AP-1/2β, as were other trafficking-associated components (Fig. 5B; Suppl. Table 4). However, no peptides from AP-2 α, µ, and σ subunits were identified in MS analysis of four replicate pulldowns (Fig. 5B, 5C). AP-2 independent clathrin-dependent endocytosis has been demonstrated in African trypanosomes, where the genes encoding the AP-2 subunits are absent, and also suggested in *T. cruzi* where clathrin does not appear to interact with AP-2, but has never before been demonstrated in *Plasmodium*. AP-3 complex subunits were also identified but were not enriched compared to controls. The role of the AP-4 complex is unclear in eukaryotes, but this complex is not believed to interact with clathrin. Consistent with this, we did not detect peptides corresponding to the *P. falciparum* AP-4 complex. Interestingly, we found peptides corresponding to Sortilin, Vps9, a DnaJ chaperone, and RESA enriched in the clathrin interactome, among other exported and trafficking-related factors. Collectively these data suggest that retromer, *trans*-Golgi, and secretory trafficking likely involve clathrin-coated vesicles (Suppl. Table 4). Consistent with these diverse active processes, our localisation of PfCHC demonstrates many rapidly-cycling foci decorating the plasma membrane and cytoplasmic structures in live trophozoites (Fig. 5F). This dynamic localisation was never observed for AP-2 in our imaging studies. These data support a novel, clathrin-independent role for the AP-2 complex in *Plasmodium*.

### Disruption of trafficking confers high-level artemisinin resistance *in vitro*

As a first step to explore the mechanism of the AP-2µ(I592T)-mediated artemisinin resistance phenotype reported above, we lastly proposed that disruption of intracellular traffic would increase parasite tolerance to artemisinin exposure. In higher-order eukaryotes, blockade of secretory traffic instigates an eIF2α-dependent cell stress response ^44^, a process recently shown to underlie K13-mediated artemisinin resistance in *P. falciparum* ^45^. Interestingly, as transient BFA exposure rapidly and reversibly inhibits secretory traffic from the ER, an impact also seen with transient incubations of parasites at 15-20°C ^46^, a recent study by our laboratory showed that transient incubations at these temperatures dramatically increased parasite survival to artemisinin during an RSA^4h^ pulse ^33^. Thus, to provide further evidence that intact secretory traffic in early ring-stage parasites is important for artemisinin susceptibility, we treated highly synchronised ring stage parasites with BFA for 1 hour from hours 2-3 post-invasion ^34^. At hour 3, we initiated the RSA^4h^ as described in the presence of this BFA, and at hour 7 we washed both compounds off extensively, returning the treated cells to culture. We compared BFA^5h^-treated 3D7 parasite survival in the RSA^4h^ to that of untreated cells (Fig. 6A). In the absence of DHA, the BFA^5h^ treatment had little effect on parasite proliferation (Fig. 6B), as previously reported, but this treatment markedly increased eIF2α-P, the cell stress indicator previously detected in artemisinin-resistant K13^mut^ parasites (Fig. 6C). In the RSA^4h^, the presence of BFA provided substantial protection, with 45% of BFA^5h^-treated parasites surviving the DHA exposure, compared with less than 3% of solvent-treated control (Fig. 6C). It is notable that we also detected low levels of eIF2α-P in solvent-treated ring stage parasites, in contrast to previous reports ^45^. These data show that transient chemical blockade of secretory traffic is sufficient to generate a K13-like artemisinin resistance phenotype, adding to the evidence that both intracellular traffic and the cell stress response may be important determinants of ring-stage parasite susceptibility to artemisinin *in vitro*.

## Discussion

In this study, we implicate the µ subunit of the *P. falciparum* AP-2 adaptor complex in artemisinin susceptibility *in vitro* and provide functional insights into the role of this complex in *Plasmodium*, highlighting several divergences from eukaryotic dogma. Specifically, we introduced a mutation, first identified in artemisinin-resistant *P. chabaudi*, into the orthologous *P. falciparum* locus by genome editing. This demonstrated an enhanced ring-stage artemisinin survival phenotype similar to, though less severe than, the genetically-distinct K13-mediated phenotypes reported in Southeast Asian strains. Interestingly, a mutation in the α-subunit of the same AP-2 complex was recently identified in an experimentally evolved artemisinin-resistant *P. falciparum* clone ^47^, providing further support for our conclusion that the AP-2 complex is capable of, and central to, modulating parasite susceptibility to artemisinin. Further, Demas *et al.* ^14^ have recently demonstrated K13-independent ring-stage resistance *in vitro* mediated by variants of the *P. falciparum* Coronin protein. Coronin has been proposed to interact with EPS15, an AP-2 partner described in other eukaryotes, a homologue of which was identified here as interacting with AP-2μ in *P. falciparum* (Fig. 4C; Suppl. Table 3). We also clarify that the PfAP-2µ(S160N) mutation currently circulating in African parasite populations does not impact ring-stage artemisinin sensitivity in the 3D7 genetic background.

By interrogating the localisation of epitope-tagged AP-2µ using immunofluorescence and immunoelectron microscopy, we provide evidence that the AP-2 complex is localised to the plasma membrane as well as distinct cytoplasmic foci, likely corresponding to vesicles throughout the cytosol during intra-erythrocytic development. These foci segregate into merozoites during schizogony before being carried into the nascent parasite. By immunoelectron microscopy, we show that these vesicles partially co-label with ER-resident PDI, as well as with Rab5B, a protein associated with the parasite plasma membrane and digestive vacuole as trafficking destinations. Surprisingly, these vesicles also correspond well with K13-labelled structures within the resolution limits of light microscopy, though we find no evidence of direct interaction between the two molecules. Instead, AP-2µ associates with another kelch-type protein, *P. falciparum* K10, and also with potential trafficking chaperones and regulators.

We demonstrate that the AP-2 complex contains α2, μ2 and σ2 subunits plus a β-subunit that supplies both the AP-1 and AP-2 complexes. This promiscuous behaviour has been reported previously in other organisms, where it has functional importance for targeting the vesicular complex to specific membranes ^48^. Further work is required to determine if this is also true of β1/2 in *Plasmodium*. Additionally, we provide the first description of CHC in *P. falciparum* and find no evidence of interaction or co-localisation with AP-2, despite identifying other adaptor complex subunits in a CHC pulldown. This was surprising because AP-2 is the canonical clathrin adaptin complex of eukaryotes, although a similar result was obtained for AP-2 in *T. cruzi* ^43^. A previous study of AP-1µ in *P. falciparum* identified CHC as an interacting partner ^46^, and we recapitulate this, showing that all four components of the AP-1 complex immunoprecipitate with CHC. This suggests that despite the shared β subunit, which canonically links the AP complex to clathrin, other subunits or factors may be involved in coat protein selection in *Plasmodium falciparum*. The coat protein (or proteins) associated with PfAP-2 vesicles remains unknown but may lie among the many proteins of unknown function identified in our AP-2 interactome (Fig. 4; Suppl. Table 2). Future studies should aim to further define AP-2-and clathrin-mediated traffic in Apicomplexans and establish whether other adaptins perform similarly diverged non-canonical roles, potentially with impacts on drug susceptibility, immune evasion and the basic biology of the parsite cell. It remains to be determined if such AP-2-independent clathrin-trafficking is more widespread, as the presence of such a pathway in two evolutionarily distinct protsists suggests this may be a more general phenomenon.

Further to these unexpected findings, we show conditional AP-2µ deletion causes profound defects in membrane segregation and organellar integrity during schizont maturation, and prevents intra-erythrocytic growth of *P. falciparum in vitro*. It is difficult to tease out which aspects of the complex observed phenotype are secondary to AP-2µ knockout and which are primary. Co-localisation of µ2 and Rab5B by immunoelectron microscopy may suggest that the observed plasma membrane fragility, lipid droplet accumulation, and disrupted digestive vacuoles are interrelated. AP-2µ KO may disrupt retromer-like trafficking needed to recycle lipid from the digestive vacuole back to the plasma membrane, leading to lipid accumulation near the digestive vacuole and insufficient fatty acids available for growing and segregating membranes during schizogony. This potential role for AP-2 in membrane organisation in schizonts may impact the integrity of apical secretory organelles, as several key factors are mislocalised in nascent merozoites upon AP-2µ KO (Suppl. Fig. 7, 8). These observations and the specific directionality of AP-2 traffic require further investigation.

Our data, although supportive of a link between trafficking, K13, and artemisinin susceptibility, can provide no new direct insights into the elusive function of K13 in *P. falciparum*. Previous studies have suggested K13 may be a ubiquitin ligase scaffolding factor and associated with PI3P-labeled secretory vesicles in the cell ^49, 50^. A general role for K13 in intracellular traffic determined by these studies fits well with our data, but further work is necessary to reconcile those studies with others showing a conflicting role for PI3P ^51^ and implicating autophagy in modulating artemisinin susceptibility ^52^. As a recycling process, autophagy is dependent upon intracellular trafficking, and we demonstrate that AP-2 deletion clearly affects similar processes.

Interference with intracellular traffic, in our hands via either transient BFA treatment in this study or temperature modulation ^33,46^, induces a profound artemisinin resistance phenotype in genetically wildtype 3D7 parasites. As we show, BFA induces regulatory phosphorylation of eIF2α, a hallmark of the cell stress response and globally attenuated protein synthesis, which mechanistically underlies artemisinin resistance. Consistent with this, previous data indicate that similar transient BFA treatments also stall development and translation in *Plasmodium* ring stages ^34^. We interpret these data to indicate that the fidelity of intracellular trafficking is intimately linked with this stress response, and we anticipate that AP-2 mutation mechanistically disrupts the processes observed here. Further studies characterising our AP-2µ(I592T) mutant, in particular those examining membrane organisation, DV integrity, and stress response activation, are planned.

Resistance to artemisinin has long been thought to be multigenic, consistent with our model presented above ^14, 53^. So far, laboratory evolution of artemisinin resistance has mostly generated parasites with K13 mutations that are different from those most prevalent in the field, or mutations in other loci altogether, such as the recently-described *pfcoronin* ^6, 14, 47^. Similarly, the AP-2µ(I592T) mutation described in this study has not been observed in human infections. However, using the descriptive characterisation provided here of the wildtype PfAP-2µ molecule as a platform, full characterisation of the I592T variant can now proceed, with the aim of understanding the mechanism by which *in vitro* artemisinin susceptibility is significantly reduced.

## Materials and Methods

### Plasmid Design and Construction

Two plasmids were created for transfecting 3D7 parasites: pL6-AP2µ(I592T)-sgDNA and pL6-AP2µ(I592I)-sgDNA. These constructs utilized the plasmid system described by Ghorbal *et al.* and encoded the donor sequence carrying the mutation of interest and the guide RNA^54^. The *pfap2mu_*I592T donor template was created by first amplifying the 5’ homology region with primers P1 and P2 and the 3’ homology region with primers P3 and P4. These two fragments were assembled by overlap PCR using primers P1 and P4; the resulting fragment was cloned into pL6-eGFP with *Eco*RI and *Nco*I. Primers P5 and P6 were annealed to create a dsDNA molecule encoding the sgRNA sequence with 5’ and 3’ extensions to enable InFusion Cloning (Clontech) into *Btg*ZI-digested pL6. To create the AP2µ (I592I) donor fragment, primers P2 and P3 mentioned above were replaced with P7 and P8. Primer sequences are provided in Supplementary Table 22.

### Parasite Culture and Generation of Transgenic Parasites

*Plasmodium falciparum* culture was performed in AB+ red blood cells obtained from the UK National Blood & Tissue Service, in RPMI supplemented with Albumax II, L-glutamine, and gentamycin, at 5% haematocrit under 5% CO_2_ conditions at 37°C, and synchronised using 70% Percoll gradients to capture schizonts or 5% sorbitol to isolate ring-stage parasites.

To establish the techniques in our laboratory, two transfection methods were used in this study. For integration of SNPs, ring stage transfection was performed. Briefly, 3D7 parasites were cultured to approximately 10-15% parasitaemia in 5% haematocrit under standard conditions. Immediately before transfection, 100 µg of each plasmid (pL6 and pUF1-Cas9) was ethanol-acetate precipitated and resuspended in 100 µl of sterile TE. 300 µl of infected red cells were isolated by centrifugation and equilibrated in 1 × Cytomix (120 mM KCl, 5 mM MgCl_2_, 25 mM HEPES, 0.15 mM CaCl_2_, 2 mM EGTA, 10 mM KH_2_PO_4_/K_2_HPO_4_, pH 7.6). 250 µl packed cells were combined with 250 µl of 1 × Cytomix in a 2 mm transfection cuvette (Bio-Rad Laboratories). The precipitated and resuspended DNA was added to the cell suspension in the cuvette. The cells were immediately pulsed at 310V, 950 µF, and infinite resistance in a Bio-Rad Gene Pulser. The electroporated cells were then washed twice in complete media to remove debris and returned to culture. Fresh red blood cells were added to approximately 5% total haematocrit on day 1 after transfection along with 2.5 nM WR99210 and 1.5 µM DSM-1. Media and selection drugs were replenished every day for 14 days and then every 3 days until parasites were observed by microscopy. Parasites recovered approximately 3 weeks post-transfection. The tagged µ2 parasite line was created by the spontaneous DNA uptake method exactly as described by Deitsch *et al* ^55^. The 3D7-µ2-2xFKBP-GFP and 3D7-CHC-2xFKBP-GFP parasite lines were generously provided by Tobias Spielmann and generated via selection-linked integration ^36^. The 3D7 DiCre-expressing parasite line ^37^ and Cam3.II family parasite lines ^13^ were generously provided by Michael Blackman and David Fidock, respectively.

### Drug Susceptibility Assays

Synchronised ring stage parasites at 0.5% parasitaemia and 2% haematocrit were exposed to a range of dihydroartemisinin, quinine, chloroquine, lumefantrine, piperaquine, and mefloquine concentrations over a full lifecycle in two biological replicates within 96-well microtitre plates, following standard procedures in our laboratory. Serial two-fold drug dilutions were performed across the plate using complete media; parasites were added thereafter. The plate was incubated at 37°C for 48 hours, freeze-thawed, and drug sensitivity quantitated using SYBR Green fluorescence as described ^56^. Parasite survival was normalised between parasites exposed to 10 µM chloroquine and a drug-free control. EC50 estimates were determined using a non-linear regression fit in GraphPad Prism (GraphPad Software, San Diego California USA, version 7.01).

DHA drug-pulse experiments were carried out using our previously published RSA^0-4h^ format,^44^ itself a modification of the RSA^0-6h^ of Witkowski *et al.* ^27^ and the RSA^0-3h^ approach of Yang *et al.* ^32^. Survival estimates were based on FACS counts performed on live cells using 1:10,000 MitoTracker Deep Red and 1:1,000 SYBR Green (Invitrogen Molecular Probes) stain in PBS. Cells were stained using 8 volumes of stain solution per volume of culture (2% HCT) and analysed on a LSR-II Flow Cytometer (BD Biosciences). Statistical significance was determined by the two-sample Wilcoxon rank-sum (Mann-Whitney) test.

### Fluorescence Microscopy

To prepare slides for immunofluorescence microscopy, thin smears were first prepared on glass slides and quickly dried. Sections for staining were marked with a hydrophobic pen. Cells were fixed for 10 minutes with 4% formaldehyde in PBS, washed three times with PBS, permeabilised with 0.1% (v/v) Triton X-100 in PBS for 10 minutes, washed again, and blocked for 1 hour with 3% (w/v) BSA in PBS. Primary antibodies were diluted in PBS containing 3% (w/v) BSA and 0.1% (v/v) Tween20 and incubated on the slide overnight at 4°C. Slides were again washed several times with PBS and incubated with secondary antibodies diluted in the same buffer. Slides were incubated for 1 hour at room temperature and washed. Glass coverslips were mounted with 1 μl of Vectashield with DAPI. Images were taken on a Nikon TE-100 inverted microscope.

### Electron Microscopy

For ultrastructural analysis, *P. falciparum* were cultured at 37°C in 4 mL volumes in 6-well tissue culture dishes (Techno Plastic Products) at 2% hematocrit until reaching 6-10% parasitemia. Cultures were synchronized until >80% of parasites were in ring stage growth and then treated for 1 h with 10 nM rapamycin to excise *pfap2μ*. Cultures treated with DMSO were used as negative controls. Parasites were then washed twice with RPMI and incubated at 37°C until harvesting. Synchronised parasites were magnetically sorted in both the trophozoite and the schizont stages by passage through a magnetically mounted MACS LD separation column (Miltenyi Biotech, Germany), collected by centrifugation and fixed in 2% paraformaldehyde/2.5% glutaraldehyde (Polysciences Inc., Warrington, PA) in 100 mM cacodylate buffer, pH 7.2 for 1 h at room temperature. Samples were washed in cacodylate buffer and post-fixed in 1% osmium tetroxide (Polysciences Inc.) for 1 h. Samples were then rinsed extensively in dH_2_O prior to en bloc staining with 1% aqueous uranyl acetate (Ted Pella Inc., Redding, CA) for 1 h. Following several rinses in dH_2_O, samples were dehydrated in a graded series of ethanol and embedded in Eponate 12 resin (Ted Pella Inc.). Sections of 90 nm were cut with a Leica Ultracut UCT ultramicrotome (Leica Microsystems Inc., Bannockburn, IL), stained with uranyl acetate and lead citrate, and viewed on a JEOL 1200 EX transmission electron microscope (JEOL USA Inc., Peabody, MA) equipped with an AMT 8 megapixel digital camera (Advanced Microscopy Techniques, Woburn, MA).

### Immunoelectron Microscopy

Parasites at 2% hematocrit and 6-8% parasitemia were magnetically sorted from uninfected RBCs and ring stage parasites via MACS LD separation columns (Miltenyi Biotech, Germany), collected by centrifugation and fixed for 1 h at 4°C in 4% paraformaldehyde (Polysciences Inc., Warrington, PA) in 100 mM PIPES/0.5 mM MgCl_2_, pH 7.2. Samples were then embedded in 10% gelatin, infiltrated overnight with 2.3 M sucrose/20% polyvinyl pyrrolidone in PIPES/MgCl_2_ at 4°C and finally trimmed, frozen in liquid nitrogen, and sectioned with a Leica Ultracut UCT7 cryo-ultramicrotome (Leica Microsystems Inc., Bannockburn, IL). 50 nm sections were blocked with 5% fetal bovine serum (FBS)/5% normal goat serum (NGS) for 30 min and subsequently incubated with primary antibody for 1 h at room temperature (anti-PDI mouse 1:100 (1D3; Enzo Life Sciences), anti-GFP rabbit 1:200 (A-11122; Life Technologies), anti-GFP mouse 1:100 (11814460001; Roche), anti-HA rabbit 1:50 (H6908; Sigma-Aldrich), anti-Rab5a rabbit 1:50 (Gordon Langsley) and anti-Rab5b rat 1:50 (Gordon Langsley). Secondary antibodies were added at 1:30 for 1 h at RT (12 nm Colloidal Gold AffiniPure Goat anti-Rabbit IgG (H+L)(111-205-144), 18 nm Colloidal Gold AffiniPure Goat anti-Rabbit IgG (H+L)(111-215-144), 12 nm Colloidal Gold AffiniPure Goat anti-Mouse IgG (H+L)(115-205-146), and 18 nm Colloidal Gold AffiniPure Goat anti-Mouse IgG+IgM (H+L)(115-215-068) (Jackson ImmunoResearch). Sections were then stained with 0.3% uranyl acetate/2% methyl cellulose, and viewed on a JEOL 1200 EX transmission electron microscope (JEOL USA Inc., Peabody, MA) equipped with an AMT 8 megapixel digital camera and AMT Image Capture Engine V602 software (Advanced Microscopy Techniques, Woburn, MA). All labeling experiments were conducted in parallel with controls omitting the primary antibody. These controls were consistently negative at the concentration of colloidal gold conjugated secondary antibodies used in these studies.

### Antibodies

Anti-HA (Roche, 3F10 clone) was obtained commercially. Anti-Clathrin Heavy Chain (rabbit) antibodies were generously donated by Frances Brodsky. We thank Gordon Langsley for making anti-Rab5B antibodies available. Anti-eIF2a antibodies were generously gifted by William Sullivan.

### Pulldown and Mass Spectrometry

For cryomilling, lysate preparation, and immunoprecipitation, *P. falciparum* cultures were grown to approximately 8% parasitaemia in 5% haematocrit in approximately 6 L of complete medium, harvested by centrifugation at trophozoite stage and lysed with 0.15% (w/v) saponin in PBS at 4°C. Parasites were harvested by centrifugation at 13K RPM for 5 minutes at 4°C and washed several times with cold PBS to remove haemoglobin and red cell debris. The washed, packed parasites were resuspended to 50% density in PBS, flash frozen in liquid nitrogen, and stored at −80°C. This process was repeated until 6-8 ml of resuspended parasites had been stored. The frozen material was placed directly into the ball chamber of a liquid N_2_-cooled cryomill (Retsch) and milled under 7 cycles of 3 minutes cooling and 3 minutes milling. The milled powder was then carefully removed from the ball chamber and stored in liquid N_2_. All steps were performed at or below −80°C to prevent parasite material from thawing. 300 mg of the milled powder per replicate was lysed in buffer A (20 mM HEPES, 100 mM NaCl, 0.1% (v/v) Triton X-100, 0.1 mM TLCK, protease inhibitors (Complete Protease Inhibitor Cocktail tablet, EDTA-free, Roche), pH 7.4) or buffer B (20 mM HEPES, 250 mM sodium citrate, 0.1% (w/v) CHAPS, 1 mM MgCl_2_, 10 mM CaCl_2_, 0.1 mM TLCK, protease inhibitors, pH 7.4). Buffer B was previously optimised to immunoprecipitate clathrin heavy chain from *T. brucei* ^43^. The lysate was sonicated and clarified by centrifugation. For 3D7-AP-2µ-3xHA the soluble extract was incubated with 240 μl anti-HA magnetic beads (Pierce, ThermoFisher) for 1 hour. Beads were washed three times with lysis buffer several times, and bound material was eluted by suspending the beads in 80 μl non-reducing LDS buffer (Invitrogen) and incubating at 70 °C for 10 minutes. After removing the beads NuPAGE sample-reducing agent (ThermoFisher) was added to the supernatant.

The PfCHC-2xFKBP-GFP soluble extract in buffer B was incubated with 4μl recombinant anti-GFP nanobodies covalently coupled to surface-activated Epoxy magnetic beads (Dynabeads M270 Epoxy, ThermoFisher) for 1 hour. Beads were washed three times in buffer B and finally eluted in 80 μl LDS buffer (Invitrogen), supplemented with NuPAGE, at 70 °C for 10 minutes. The eluates were concentrated in a speed-vac to 30 μl and run approximately 1.2 cm into an SDS-PAGE gel. The respective gel region was sliced out and subjected to tryptic digest, reductive alkylation. Eluted peptides were analysed by liquid chromatography tandem mass spectrometry (LC-MS/MS) on a Dionex UltiMate 3000 RSLCnano System (Thermo Scientific, Waltham, MA, USA) coupled to an Orbitrap Q Exactive mass spectrometer (Thermo Scientific) at the University of Dundee Finger-Prints Proteomics facility. Mass spectra were processed using MaxQuant version 1.5, using the intensity-based label-free quantification (LFQ) method ^57, 58^. Minimum peptide length was set at six amino acids and false discovery rates (FDR) of 0.01 were calculated at the levels of peptides, proteins and modification sites based on the number of hits against the reversed sequence database. Ratios were calculated from label-free quantification intensities using only peptides that could be uniquely mapped to a given protein. The software Perseus was used for statistical analysis of the LFQ data ^59^.

### Disruption of Secretory Traffic with Brefeldin A

Synchronous wild-type 3D7 cultures were treated with 5 µg/ml BFA or solvent only from hour 2 post-invasion, and a subsequent RSA^4h^ with 700 nM DHA performed from hours 3 to 7, after which both drug and BFA were washed off, and the parasites returned to drug-free culture to a total of 72 hours post-invasion.

## Author Contributions

RCH conceived, designed, and executed the study. RCH performed cell culture, transfections, cryo-milling and pulldowns, MS sample preparation, fluorescence microscopy, and drug susceptibility assays. RLE and AOJ designed and performed electron microscopy experiments. MZ performed cryo-milling, pulldowns, and mass spectrometry analysis, supported by MCF. DVS performed cell culture and drug susceptibility assays. FM performed cell culture and assisted the design and execution of the study. MH and RM assisted the design and execution of the study. SDN performed cell culture and assisted the design and execution of the study. AP and CF supported the design and execution of the study. DB assisted with study design and supported the study. CJS conceived, designed and supported the study, and performed statistical analyses. RCH and CJS wrote the manuscript, with input from all other authors.

## Acknowledgements

We would like to extend our gratitude to members of the Departments of Infection and Immunity and Pathogen Molecular Biology at LSHTM for their mentorship and helpful conversations. We thank Wandy Beatty of the Washington University Molecular Microbiology Imaging Facility for helpful assistance. We thank Douglas Lamond and the Fingerprints Proteomics facility at the University of Dundee for invaluable support. RCH was supported by the UK Foreign and Commonwealth Office through the Marshall Scholarship Programme; CJS is supported by Public Health England; MZ and MCF are supported by Wellcome Trust 204697/Z/16/Z (to MCF). MCF is a Wellcome Trust Investigator. AOJ is supported by National Institutes of Health R01 AI103280 and R01 AI123433, and is a Burroughs Wellcome Fund Investigator in the Pathogenesis of Infectious Diseases (PATH). RWM and FM are supported by an MRC Career Development Award (MR/M021157/1) jointly funded by the UK Medical Research Council and Department for International Development. MNH is supported by a Bloomsbury Colleges research studentship. The Pf-Rab5b antibody was kindly provided by Gordon Langsley (Institut Cochin, France).

## Competing Financial Interests

The authors declare no competing financial interests.

## SUPPLEMENTARY INFORMATION

**Suppl. Fig. 1.**
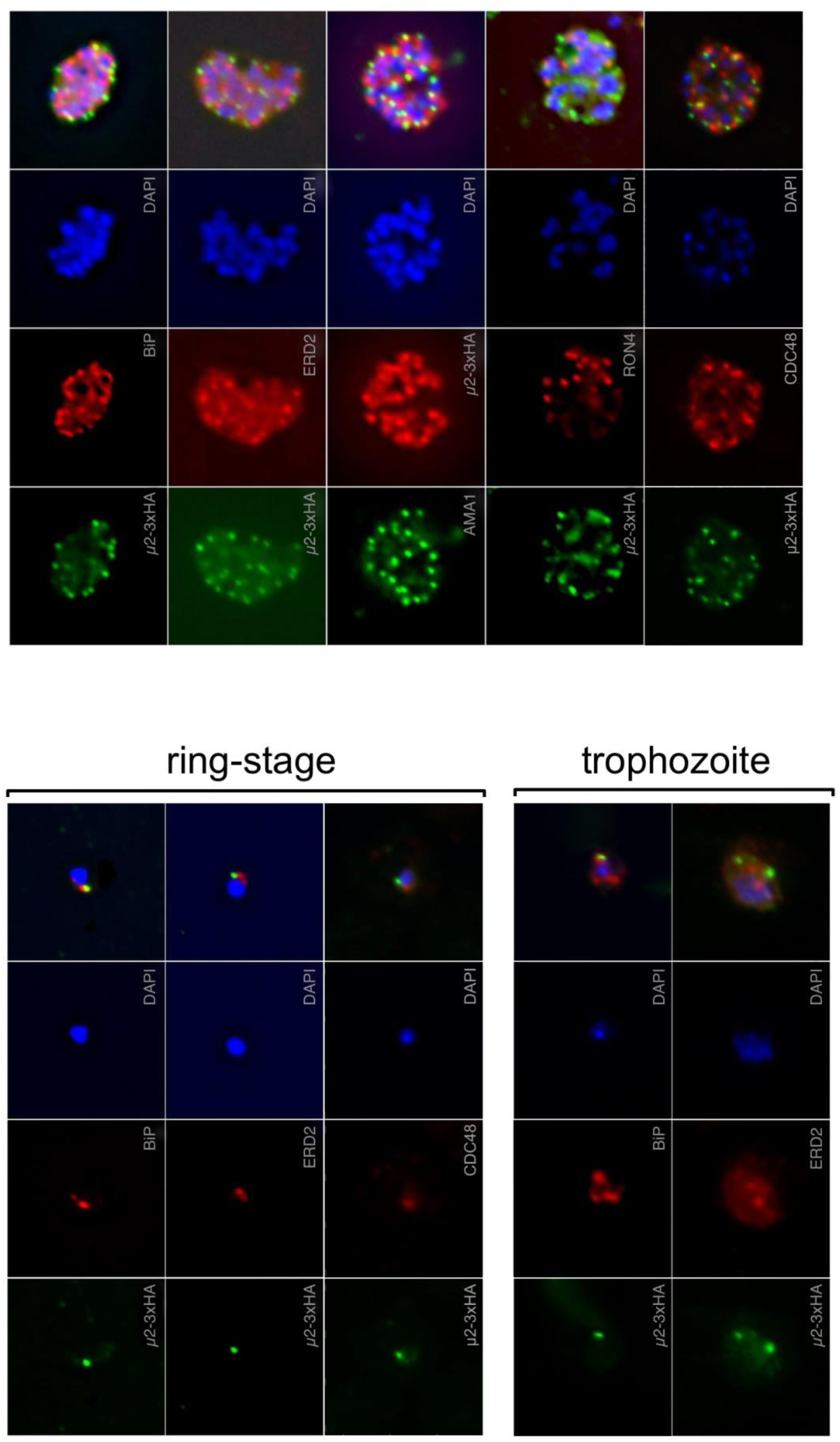
Localisation of AP-2µ-3xHA with respect to cellular landmarks.

**Suppl. Fig. 2.**
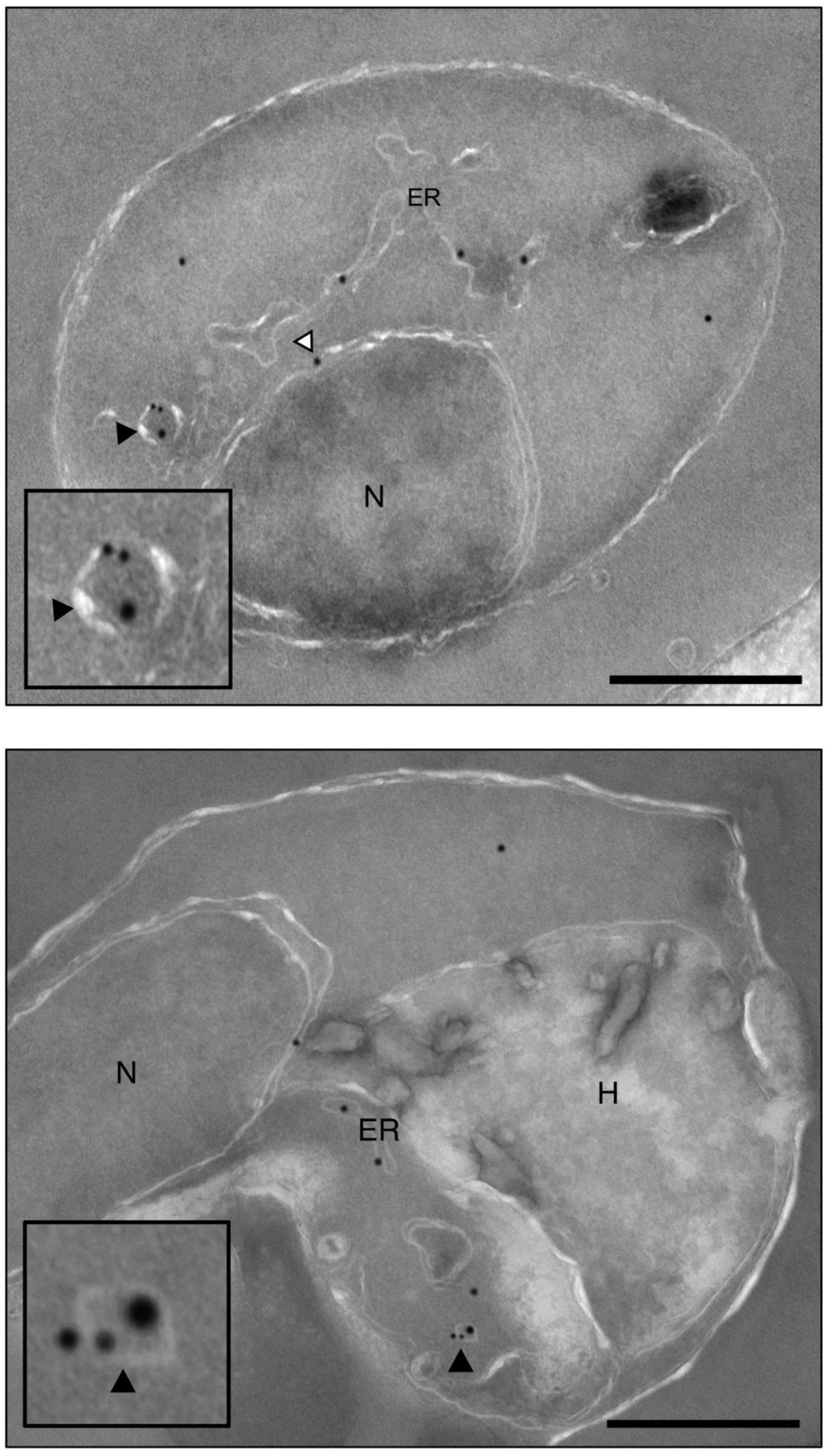
Localisation of AP-2µ-3xHA and Rab5B by immunoelectron microscopy. Electron micrographs of vesicular colocalisation of AP-2µ-3xHA (18 nm gold particles) and Rab5B (12 nm gold particles) in developing trophozoites. N: nucleus; ER: endoplasmic reticulum; H: haemozoin; white arrowhead: AP-2µ at the nuclear membrane; black arrowhead: vesicles co-labelled for both proteins. Scale bar: 500 nm.

**Suppl. Fig. 3.**
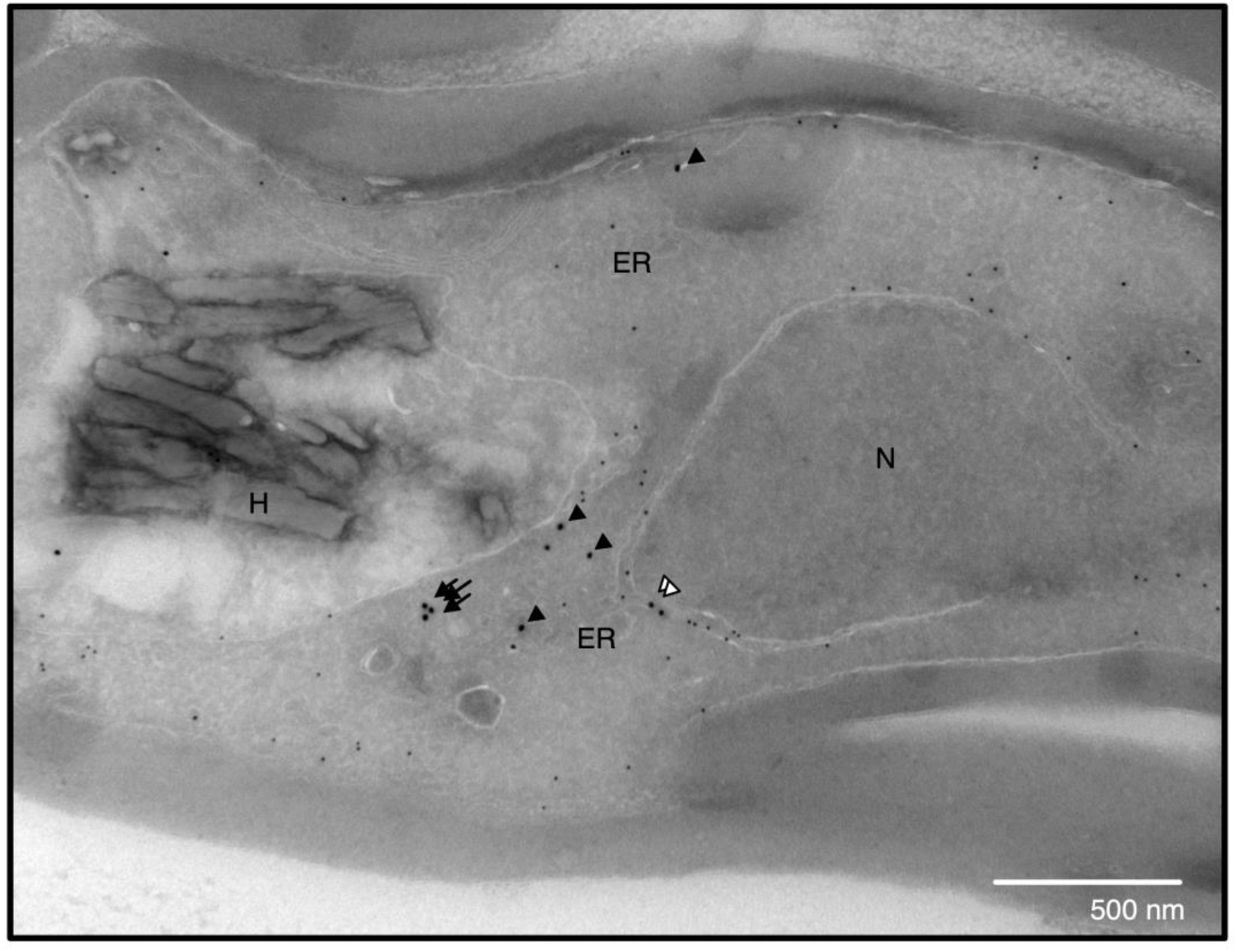
Localisation of AP-2µ-2xFKBPP-GFP and PDI by immunoelectron microscopy. Electron micrographs of vesicular colocalisation of AP-2µ-2xFKBP-GFP (18 nm gold particles) and PDI (12 nm gold particles) in developing trophozoites. N: nucleus; ER: endoplasmic reticulum; H: haemozoin; white arrowhead: AP-2µ at the nuclear membrane; black arrow: AP-2µ in vesicles; black arrowhead: AP-2µ at the ER. Scale bar: 500 nm.

**Suppl. Fig. 4.**
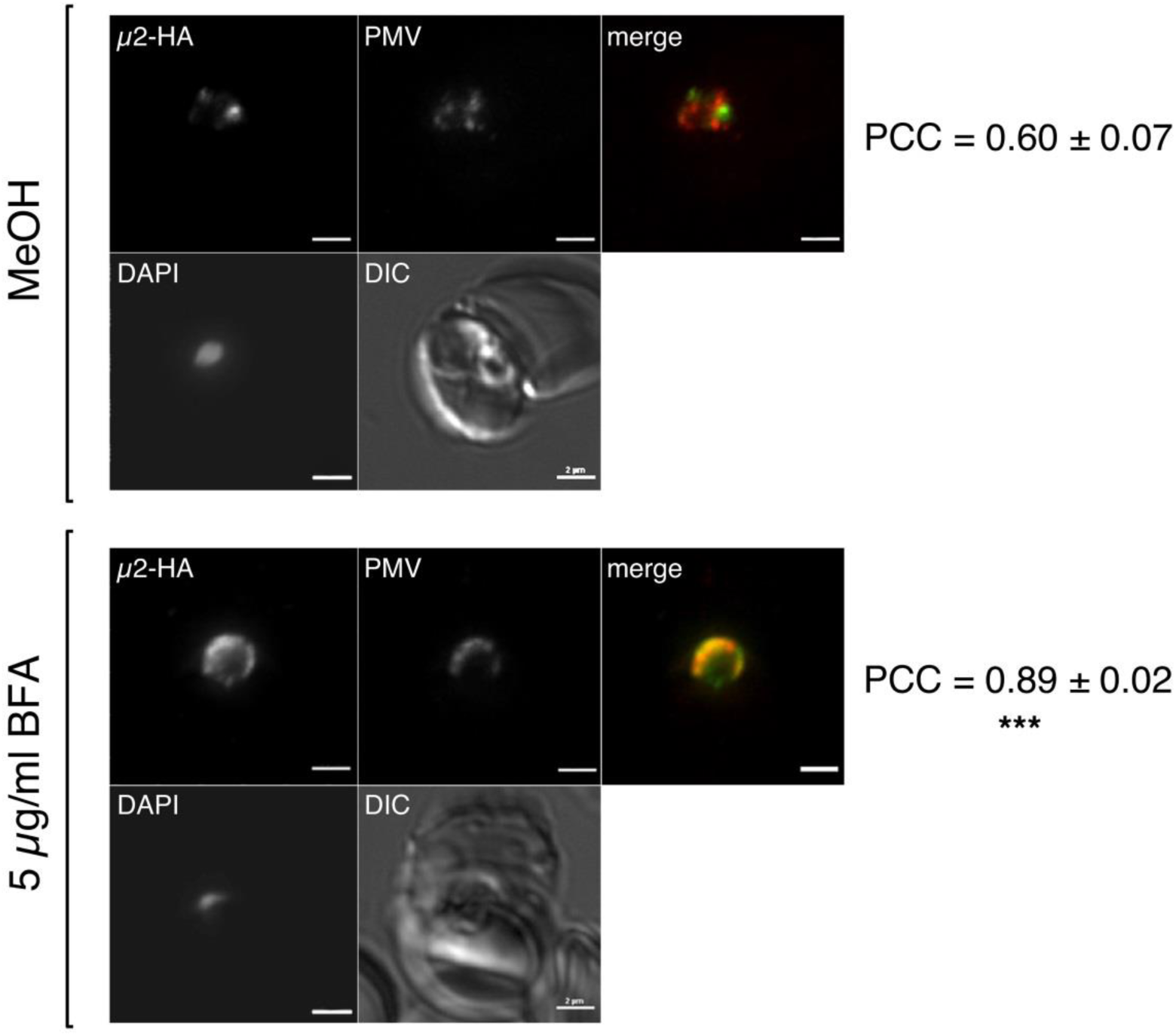
Impact of BFA on localisation of AP-2µ-3xHA. Treatment of synchronised ring-stage parasite cultures with 5 µg/ml BFA or equivalent methanol solvent for 16h and immune-stained for AP-2µ (false-coloured green) and plasmepsin V (PMV; red). Cells were fixed and stained in suspension and mounted onto coverslips. Pearson’s Correlation Coefficients (PCC) between indirect AP-2µ and PMV signals were calculated using Nikon AR Analysis software. Mean of at least 20 cells with standard deviation is shown. PCC were significantly different (***, P < 0.005) using a Student’s *t-*test. Scale: 2 µm.

**Suppl. Fig. 5.**
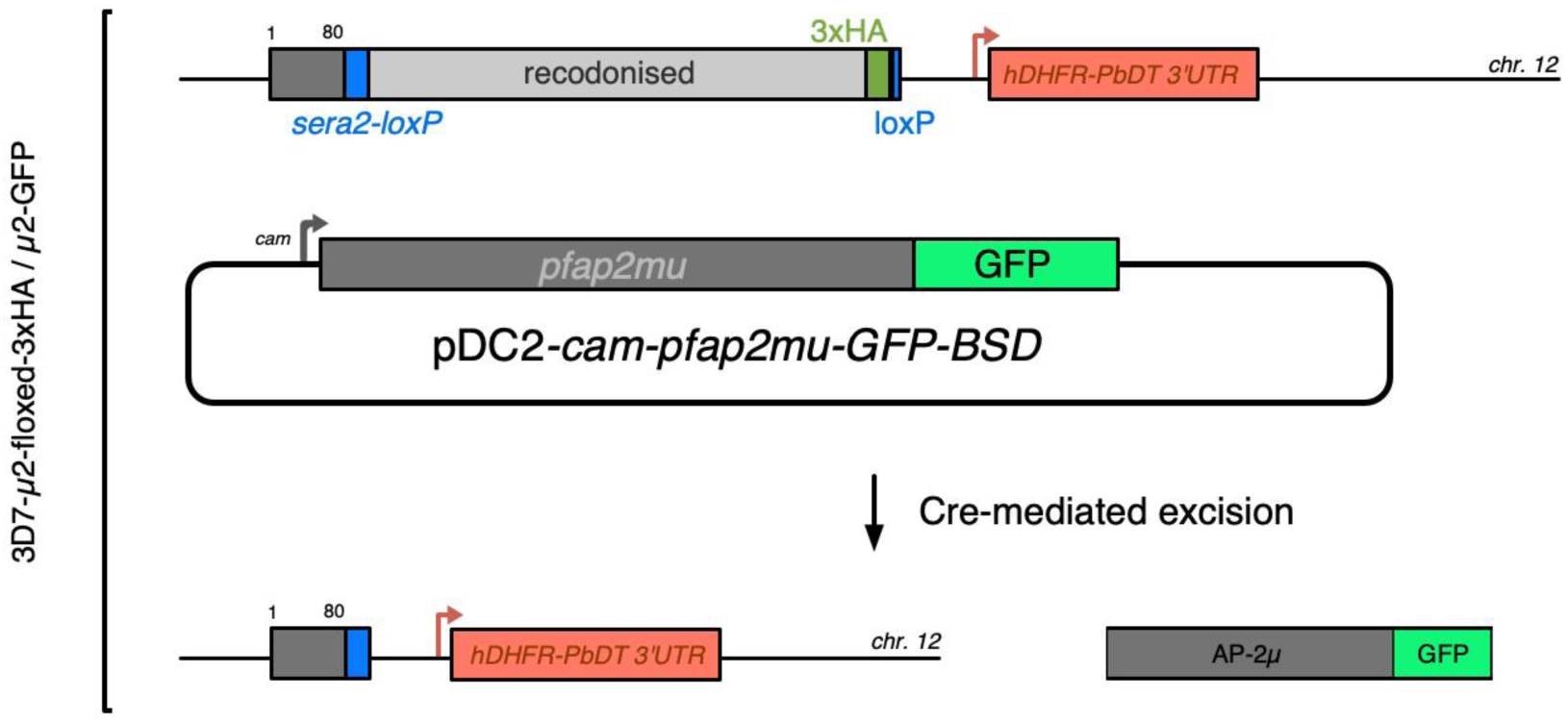
Schematic of episomal complementation of AP-2µ knockout. The *floxed* AP-2µ-3xHA-and DiCre-expressing parasite line 3D7-µ2-floxed-3xHA was transfected with a plasmid that constitutively expresses µ2-GFP under the *cam* promoter (pDC2-*cam-pfap2mu-GFP-BSD*). These cells were maintained on 2.5 µg/ml blasticidin-S (BSD). Upon addition of rapamycin, split Cre recombinase dimerises and excises the endogenous *pfap2mu* locus on chromosome 12. Parasites still express µ2-GFP via the pDC2 episome.

**Suppl. Fig. 6.**
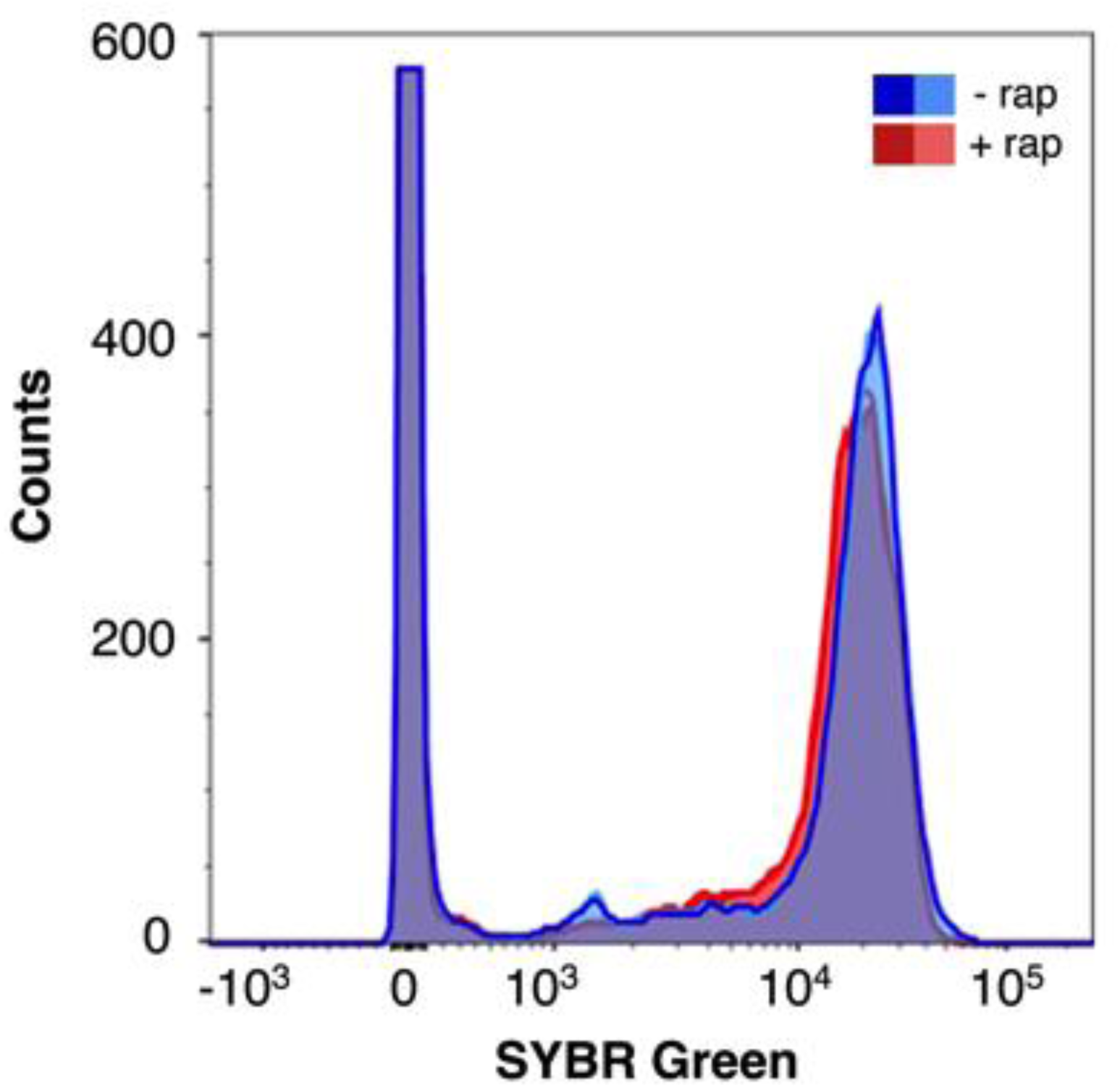
AP-2µ-KO schizonts have similar nucleic acid content to wild-type schizonts. Representative histogram comparing DNA/RNA content in wild-type (−rap, blue) and AP-2µ-KO (+rap, red) schizonts. Nucleic acid was stained using SYBR Green, and SYBR Green signal was detected in live cells by FACS. Rap-parasites were egress-blocked by treatment with the reversible PKG inhibitor Compound 2.

**Suppl. Fig. 7.**
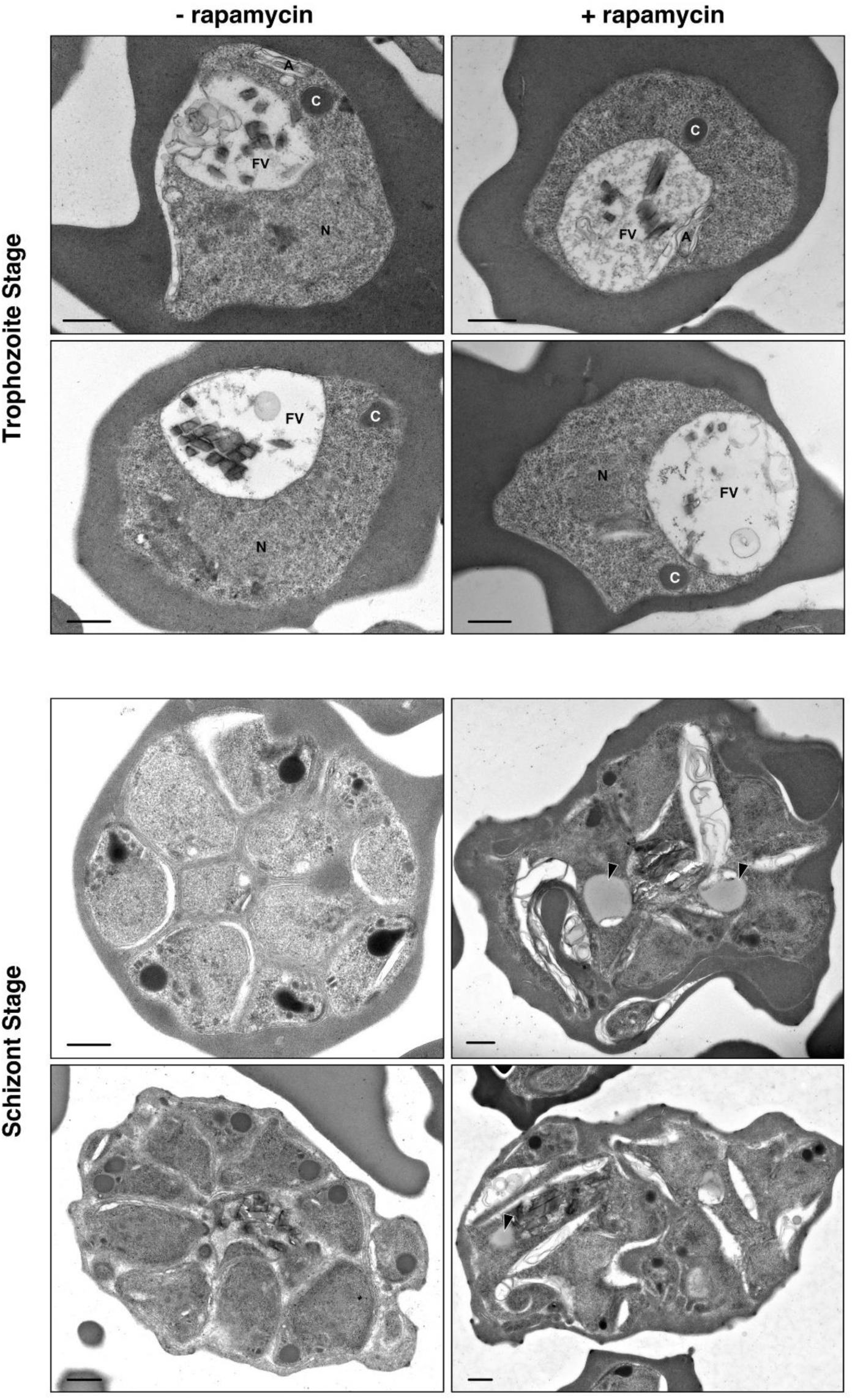
Impact of AP-2µ KO on trophozoite and schizont maturation and morphology. Electron micrographs examining the morphology of wild-type (−rapamycin) and AP-2µ-KO (+rapamycin) trophozoites (top panel) and schizonts (bottom panel). Two representative micrographs are shown for each cell type. Rapamycin or equivalent DMSO was added to synchronous ring-stage parasites for 1h and parasites were fixed for imaging in the same development cycle. 300 schizonts from each treatment were systematically enumerated for key features (see main text). Black arrowheads indicate accumulated lipid bodies in AP-2µ-KO schizonts. Labels used in trophozoite micrographs N: nucleus; FV: food (digestive) vacuole; C: cytostome; A: apicoplast.

**Suppl. Fig. 8.**
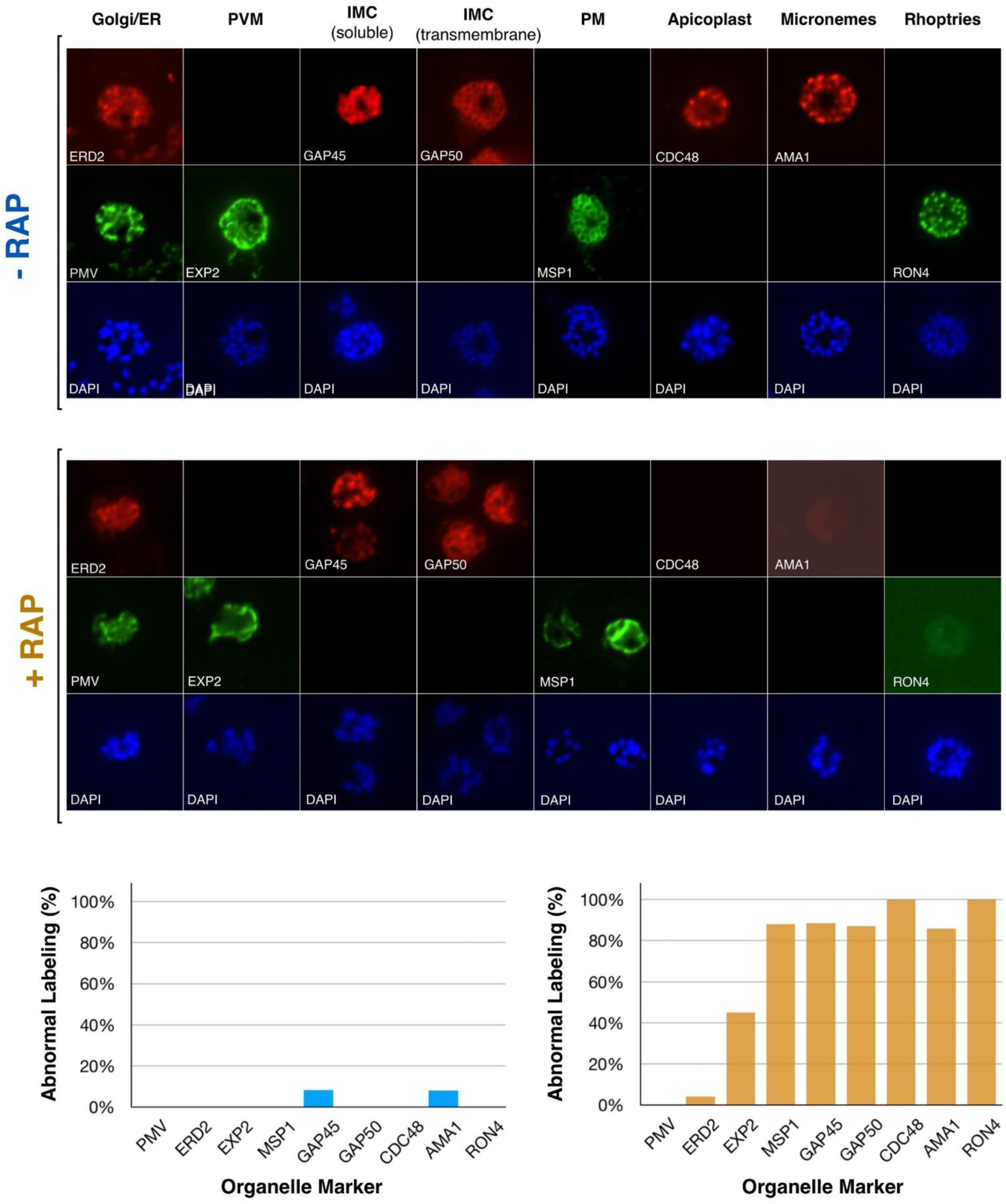
AP-2µ-KO severely disrupts schizont maturation. Antibodies against the Golgi apparatus (ERD2), ER (PMV), PVM (EXP2), IMC (GAP45, GAP50), PM (MSP1), apicoplast (CDC48), micronemes (AMA1), and rhoptries (RON4) were used to stain egress-blocked wild type (−RAP) or µ2-KO (+RAP) schizonts. Top panel shows representative IFAs for each organelle marker. Abnormal labelling was quantitated relative to the staining observed in the majority of wild type schizonts (lower panel; left, rap-; right, rap+): ERD2, PMV, CDC48: discrete punctate staining corresponding to each nucleus; EXP2: contiguous, circular, peripheral membrane staining; GAP45, GAP50, MSP1: distinct, circular grape-like staining surrounding each daughter nucleus; AMA1, RON4: discrete, apical punctate spots corresponding to each nucleus. At least 100 cells were scored for each marker. IMC: inner membrane complex; PM: plasma membrane; PVM: parasitophorous vacuolar membrane

**Suppl. Fig. 9.**
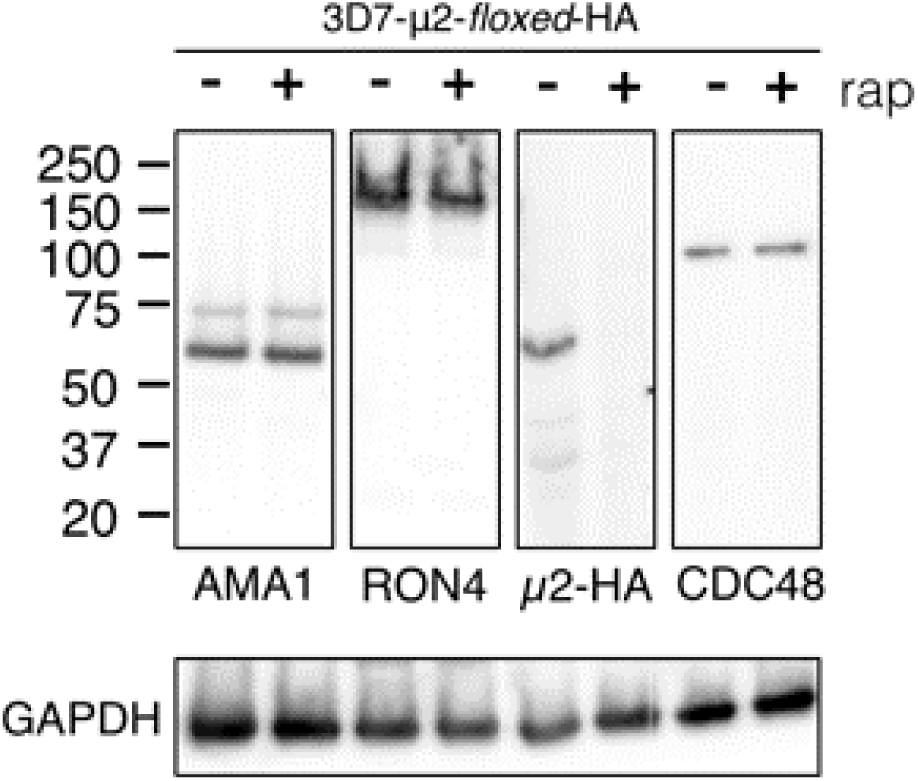
Organelle markers are mislocalised by AP-2µ KO. Antibodies against RON4, AMA1, and CDC48, markers of the rhoptries, micronemes, and apicoplast, failed to stain schizonts lacking µ2 (Suppl. Figure 4). Western blot analysis of whole-cell lysates prepared from wild type and µ2-KO schizonts confirms the overall cellular abundance of these factors is not affected by µ2-KO.

**Suppl. Table 1.**
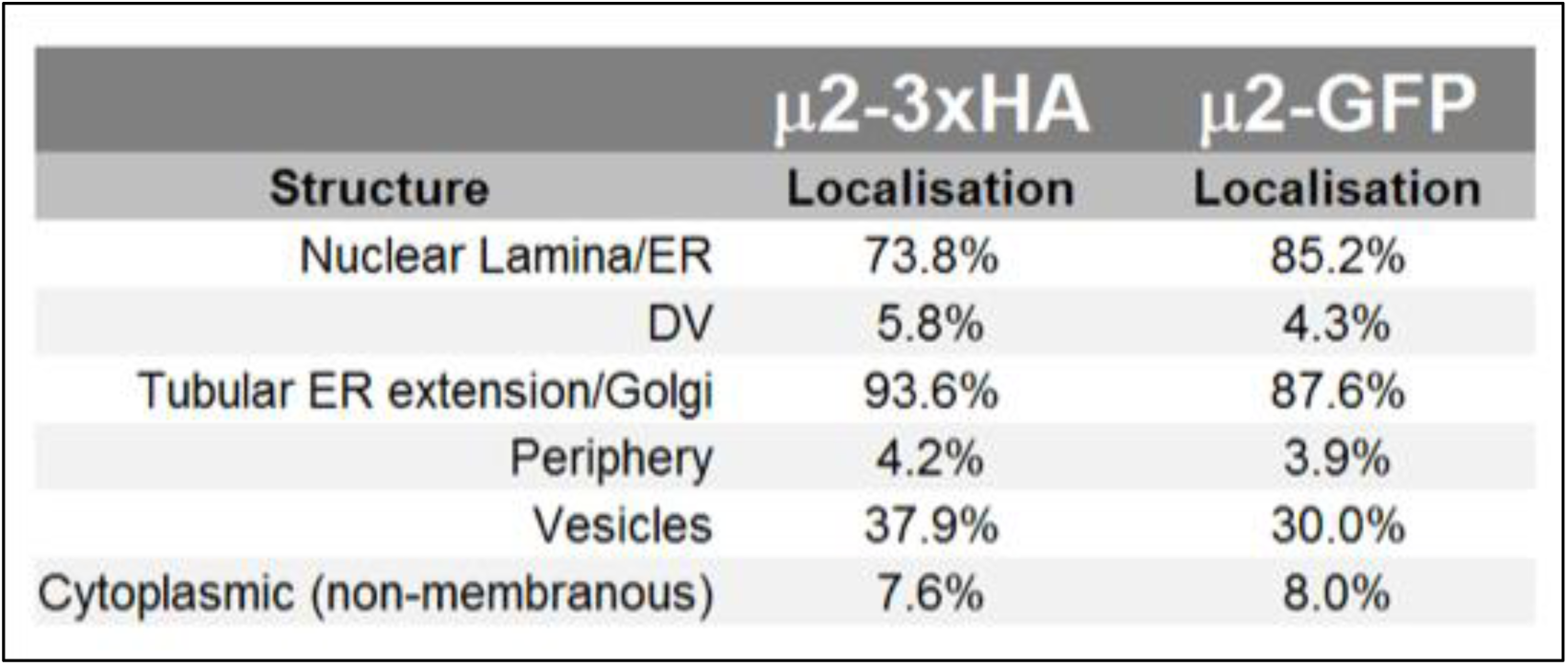
Quantitation of distribution of gold labels in immunoelectron micrographs.

**Suppl. Table 2.**
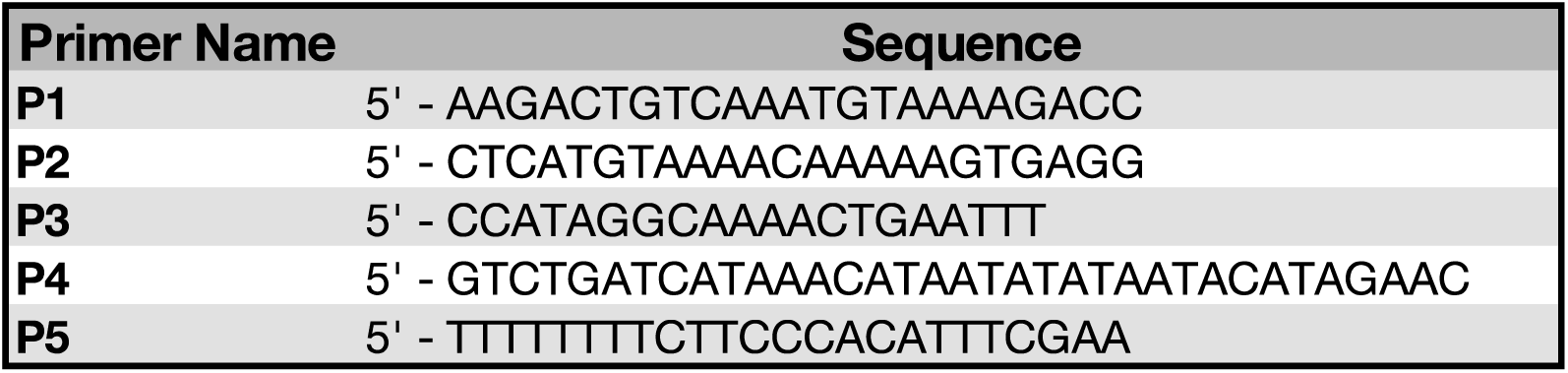
Primers used to screen transgenic lines described in this study.

**Supplementary Table 3.** Extended IP-MS interactome of AP-2µ. Factors identified with greater than 3-fold enrichment in HA pulldown in either condition compared to control pulldowns with statistical difference between HA and control pulldowns greater than 0 and at least 2 peptides per protein. ∞ indicates that no peptides were detected in the control pulldown; ND indicates protein was not identified in either the control or HA pulldown.

| Protein ID | Protein Name | Average-Enrich.<br>Triton | Average-Enrich.<br>CHAPS |
| --- | --- | --- | --- |
| PF3D7_1147300 | Conserved Plasmodium protein, unknown function | ∞ | ND |
| PF3D7_1459000 | ATP-dependent RNA helicase DBP5 | ∞ | ND |
| PF3D7_1251600 | Conserved Plasmodium protein, unknown function | ∞ | ND |
| PF3D7_0720100 | Small subunit rRNA processing protein, putative | ∞ | ND |
| PF3D7_1123400 | Translation elongation factor EF-1, subunit a | ∞ | ND |
| PF3D7_1027400 | DNA-directed RNA polymerase II subunit RPB7 | ∞ | ND |
| PF3D7_0208700 | Conserved Plasmodium protein, unknown function | ∞ | ND |
| PF3D7_1471400 | Diacylglycerol kinase | ∞ | ND |
| PF3D7_0611700 | 60S ribosomal protein L39 | ∞ | ND |
| PF3D7_1138800 | WD repeat-containing protein | ∞ | ∞ |
| PF3D7_1434300 | Hsp70/Hsp90 organizing protein | ∞ | ∞ |
| **PF3D7_1218300** | **AP-2 complex subunit mu** | 28.65 | 341.39 |
| PF3D7_1114200 | GTPase-activating protein, putative | 21.83 | ∞ |
| PF3D7_0617100 | AP-2 complex subunit alpha | 21.01 | 240.76 |
| PF3D7_1022600 | Kelch protein K10 | 19.82 | 0.07 |
| PF3D7_0217300 | AP-2 complex subunit sigma | 17.10 | ∞ |
| PF3D7_1461300 | 40S ribosomal protein S28e | 9.75 | ND |
| PF3D7_0710400 | DNA repair protein RAD14 | 8.44 | ND |
| PF3D7_0802000 | Glutamate dehydrogenase | 8.33 | 5.06 |
| PF3D7_1137400 | UVB-resistance protein UVR8 homologue | 7.54 | ND |
| PF3D7_1205900 | Conserved protein, unknown function | 7.45 | ∞ |
| PF3D7_1402500 | Ribosomal protein S27a | 6.60 | ∞ |
| PF3D7_1470700 | Conserved Plasmodium protein | 6.60 | ND |
| PF3D7_1230900 | Serine/threonine protein kinase RIO1 | 6.30 | ∞ |
| PF3D7_0508600 | Conserved Plasmodium protein, unknown function | 6.16 | ND |
| PF3D7_0321100 | Conserved Plasmodium protein, unknown function | 5.90 | ND |
| PF3D7_0813000 | Conserved Plasmodium protein, unknown function | 5.19 | ND |
| PF3D7_0801800 | Mannose-6-phosphate isomerase | 5.15 | ND |
| PF3D7_0529500 | Cell cycle regulator protein | 5.07 | ND |
| PF3D7_0212300 | Peptide chain release factor subunit 1 | 4.93 | ND |
| PF3D7_0205800 | PH domain-containing protein | 4.74 | ND |
| PF3D7_0102200 | Ring-infected erythrocyte surface antigen | 4.66 | ND |
| PF3D7_1149200 | Ring-infected erythrocyte surface antigen | 4.62 | ND |
| PF3D7_0303300 | DNA-directed RNA polymerases I, II, and III subunit RPABC2 | 4.45 | ND |
| PF3D7_0304200 | EH domain-containing protein (EPS15) | 4.43 | ∞ |
| PF3D7_1111000 | tRNA m5C-methyltransferase | 4.23 | ND |
| PF3D7_1109400 | Essential nuclear protein 1 | 4.18 | ND |
| PF3D7_0106700 | Small ribosomal subunit assembling AARP2 protein | 4.18 | ND |
| PF3D7_1404500 | rRNA biogenesis protein RRP5 | 4.09 | ND |
| PF3D7_1205700 | Targeted glyoxalase II | 3.90 | ND |
| PF3D7_1104400 | Thioredoxin | 3.85 | ND |
| PF3D7_1451900 | Ribosome biogenesis protein TSR1 | 3.83 | ∞ |
| PF3D7_0731300 | Plasmodium exported protein (PHISTb), unknown function | 3.83 | ND |
| PF3D7_0813600 | Translation initiation factor SUI1 | 3.70 | ND |
| PF3D7_0416400 | Histone acetyltransferase | 3.70 | ND |
| PF3D7_0803100 | U3 small nucleolar RNA-associated protein 14 | 3.65 | ND |
| PF3D7_1471000 | RNA 3-terminal phosphate cyclase-like protein | 3.61 | ND |
| PF3D7_0522300 | 18S rRNA (guanine-N(7))-methyltransferase | 3.54 | ND |
| PF3D7_0708500 | Heat shock protein 86 family protein | 3.52 | ND |
| PF3D7_0528100 | AP-1 complex subunit beta | 3.49 | 26.11 |
| PF3D7_1129100 | Parasitophorous vacuolar protein 1 | 3.47 | 11.66 |
| PF3D7_0722000 | Conserved Plasmodium protein, unknown function | 3.45 | ∞ |
| PF3D7_1010600 | Eukaryotic translation initiation factor 2 subunit | 3.41 | ND |
| PF3D7_0408500 | Flap endonuclease 1 | 3.41 | ∞ |
| PF3D7_1108000 | IWS1-like protein | 3.39 | ND |
| PF3D7_0106000 | Conserved Plasmodium protein | 3.30 | ND |
| PF3D7_0417800 | Cdc2-related protein kinase 1 | 3.19 | ND |
| PF3D7_0903400 | ATP-dependent RNA helicase DDX60 | 3.16 | ND |
| PF3D7_1033300 | Conserved Plasmodium protein, unknown function | 3.09 | ND |
| PF3D7_0907600 | Translation initiation factor SUI1 | 3.08 | ND |
| PF3D7_0205600 | Conserved Plasmodium protein | 3.04 | ND |
| PF3D7_1407500 | Multifunctional methyltransferase subunit TRM | 3.04 | ∞ |
| PF3D7_0926900 | Replication termination factor | 3.00 | ND |
| PF3D7_0903400 | RNA export binding protein | 1.73 | ∞ |
| PF3D7_1033300 | peroxiredoxin (oxidative stress defense) | 1.63 | 159.73 |
| PF3D7_0907600 | Parasitophorous vacuolar protein 2 | 1.30 | ∞ |
| PF3D7_0205600 | 60S Ribosomal Protein L23 | 0.83 | 24.54 |
| PF3D7_1407500 | 60S Ribosomal Protein L26 | 0.81 | 15.67 |
| PF3D7_0926900 | AP2 domain transcription factor AP2-G (nuclear) | 0.53 | 27.51 |
| PF3D7_1341300 | 60S Ribosomal Protein L18 | 0.49 | 15.35 |
| PF3D7_1237200 | Conserved Plasmodium protein, unknown function | 0.46 | 10.11 |

**Supplementary Table 4.** Extended IP-MS interactome of clathrin heavy chain. Factors identified with greater than 3-fold enrichment in GFP pulldowns compared to control pulldowns with statistical difference between GFP and control pulldowns greater than 0 and at least 2 peptides per protein. ∞ indicates that no peptides were detected in the control pulldown; ND indicates protein was not identified in either the control or HA pulldown.

| Protein ID | Protein Name | Average-LFQ<br>wt (N=3) | Average-LFQ<br>GFP (N=4) | Enrichment | -log(P) |
| --- | --- | --- | --- | --- | --- |
| PF3D7_1421000 | DIX domain-containing protein | ND | 9.42E+08 | ∞ | 3.34 |
| PF3D7_1411300 | DnaJ protein | ND | 7.14E+08 | ∞ | 5.05 |
| PF3D7_0903400 | ATP-dependent RNA helicase DDX60 | ND | 5.67E+08 | ∞ | 0.25 |
| PF3D7_API0180 | Apicoplast ribosomal protein L16 | ND | 1.30E+08 | ∞ | 0.62 |
| PF3D7_0815800 | Vacuolar protein sorting-associated protein 9 | ND | 1.25E+08 | ∞ | 2.26 |
| PF3D7_1037600 | TFIIH basal transcription factor complex helicase XPB subunit | ND | 7.32E+07 | ∞ | 0.27 |
| PF3D7_0806800 | V-type proton ATPase subunit A | ND | 6.37E+07 | ∞ | 2.09 |
| PF3D7_0202600 | Nucleic acid binding protein | ND | 6.27E+07 | ∞ | 0.40 |
| PF3D7_1210100 | Syntaxin, Qa-SNARE family | ND | 5.22E+07 | ∞ | 0.46 |
| PF3D7_0907700 | Proteasome activator 28 subunit beta | ND | 4.51E+07 | ∞ | 0.12 |
| PF3D7_1458000 | Cysteine proteinase falcipain 1 | ND | 3.84E+07 | ∞ | 0.64 |
| PF3D7_0316500 | Kinetochore protein NUF2 | ND | 3.15E+07 | ∞ | 0.01 |
| PF3D7_0301700 | Plasmodium exported protein, unknown function | ND | 3.12E+07 | ∞ | 0.49 |
| PF3D7_0314100 | Vesicle transport v-SNARE protein | ND | 2.95E+07 | ∞ | 0.60 |
| PF3D7_1006700 | Conserved Plasmodium protein, unknown function | ND | 2.94E+07 | ∞ | 0.06 |
| PF3D7_1118100 | AP-1 complex subunit sigma | ND | 2.90E+07 | ∞ | 0.15 |
| PF3D7_0802400 | Conserved Plasmodium protein, unknown function | ND | 2.89E+07 | ∞ | 0.00 |
| PF3D7_0306100 | Conserved Plasmodium protein, unknown function | ND | 2.79E+07 | ∞ | 0.21 |
| PF3D7_0820500 | Protein transport protein YIF1 | ND | 2.71E+07 | ∞ | 0.76 |
| PF3D7_0320100 | Protein transport protein SEC22 | ND | 2.51E+07 | ∞ | 0.02 |
| PF3D7_1328100 | Proteasome subunit beta type-7 | ND | 2.36E+07 | ∞ | 0.41 |
| PF3D7_1110200 | Pre-mRNA-processing factor 6 | ND | 2.21E+07 | ∞ | 0.02 |
| PF3D7_0914900 | BSD-domain protein | ND | 2.10E+07 | ∞ | 0.26 |
| PF3D7_0103200 | Nucleoside transporter 4 | ND | 1.89E+07 | ∞ | 0.48 |
| PF3D7_0529000 | Conserved Plasmodium protein, unknown function | ND | 1.86E+07 | ∞ | 0.62 |
| PF3D7_0205500 | DNA-directed RNA polymerase II 16 kDa subunit | ND | 1.86E+07 | ∞ | 0.03 |
| PF3D7_1329500 | Conserved protein, unknown function | ND | 1.58E+07 | ∞ | 0.17 |
| PF3D7_1464700 | ATP synthase (C/AC39) subunit | ND | 1.23E+07 | ∞ | 0.15 |
| PF3D7_1004400 | RNA-binding protein | ND | 1.22E+07 | ∞ | 0.37 |
| PF3D7_0931800 | Proteasome subunit beta type-6 | ND | 9.31E+06 | ∞ | 0.07 |
| PF3D7_1419400 | Conserved Plasmodium membrane protein, unknown function | ND | 9.13E+06 | ∞ | 0.12 |
| PF3D7_1435500 | Clathrin light chain | 7.17E+07 | 2.19E+09 | 30.56 | 2.74 |
| **PF3D7_1219100** | **Clathrin heavy chain** | 3.07E+09 | 6.87E+10 | 22.38 | 2.81 |
| PF3D7_1307700 | TOM1-like protein | 3.32E+07 | 6.91E+08 | 20.81 | 2.78 |
| PF3D7_1459600 | Conserved Plasmodium protein, unknown function | 1.17E+08 | 2.33E+09 | 19.89 | 2.31 |
| PF3D7_1432800 | HP12 protein homolog | 1.72E+07 | 2.14E+08 | 12.40 | 1.63 |
| PF3D7_1311400 | AP-1 complex subunit mu-1 | 2.93E+07 | 3.18E+08 | 10.85 | 2.42 |
| PF3D7_1353200 | Membrane associated histidine-rich protein | 5.61E+06 | 5.30E+07 | 9.45 | 0.35 |
| PF3D7_1016300 | GBP130 protein | 1.75E+07 | 1.56E+08 | 8.92 | 2.38 |
| PF3D7_1455500 | AP-1 complex subunit gamma | 7.43E+07 | 6.48E+08 | 8.72 | 2.27 |
| PF3D7_1308200 | Carbamoyl phosphate synthetase | 2.60E+08 | 1.71E+09 | 6.57 | 3.07 |
| PF3D7_1451800 | Sortilin | 6.96E+08 | 4.47E+09 | 6.42 | 1.51 |
| PF3D7_1408700 | Conserved protein, unknown function | 3.24E+08 | 1.99E+09 | 6.15 | 1.33 |
| PF3D7_1214900 | Conserved protein, unknown function | 2.96E+07 | 1.71E+08 | 5.78 | 1.93 |
| PF3D7_0704400 | Phosphoinositide-binding protein | 4.90E+07 | 2.72E+08 | 5.55 | 1.58 |
| PF3D7_0102200 | Ring-infected erythrocyte surface antigen | 5.42E+07 | 2.64E+08 | 4.87 | 2.16 |
| PF3D7_0528100 | AP-1 complex subunit beta, putative \| tr | 1.25E+08 | 5.69E+08 | 4.56 | 2.06 |
| PF3D7_1215900 | Serpentine receptor | 2.48E+07 | 1.11E+08 | 4.47 | 1.27 |
| PF3D7_0530100 | SNARE protein | 1.38E+07 | 5.95E+07 | 4.32 | 1.18 |
| PF3D7_0936000 | Ring-exported protein 2 | 1.76E+07 | 6.67E+07 | 3.80 | 0.90 |
| PF3D7_0103900 | Parasite-infected erythrocyte surface protein | 2.84E+07 | 1.04E+08 | 3.65 | 1.28 |
| PF3D7_0412000 | LITAF-like zinc finger protein | 2.06E+07 | 7.26E+07 | 3.52 | 0.36 |
| PF3D7_1330400 | ER lumen protein retaining receptor 1 | 1.01E+08 | 3.52E+08 | 3.50 | 1.26 |
| PF3D7_0716300 | Conserved protein, unknown function | 1.03E+08 | 3.59E+08 | 3.47 | 1.36 |
| PF3D7_1320000 | Golgi protein 1 | 2.77E+07 | 9.49E+07 | 3.42 | 1.22 |
| PF3D7_1359600 | Conserved Plasmodium protein, unknown function | 8.12E+08 | 2.77E+09 | 3.41 | 0.35 |
| PF3D7_1015600 | Heat shock protein 60 | 3.59E+07 | 1.22E+08 | 3.38 | 0.99 |
| PF3D7_1010100 | PI31 domain-containing protein | 2.63E+07 | 8.86E+07 | 3.38 | 1.40 |
| PF3D7_0532100 | Early transcribed membrane protein 5 | 1.01E+08 | 3.36E+08 | 3.32 | 1.08 |
| PF3D7_0936800 | Plasmodium exported protein (PHISTc), unknown function | 3.30E+07 | 1.08E+08 | 3.27 | 1.12 |
| PF3D7_1123500 | Golgi protein 2 | 2.97E+07 | 9.70E+07 | 3.26 | 1.10 |
| PF3D7_0922100 | Ubiquitin-like protein | 2.32E+07 | 7.14E+07 | 3.07 | 0.17 |
| PF3D7_1239700 | ATP-dependent zinc metalloprotease FTSH 1 | 5.34E+07 | 1.60E+08 | 3.00 | 1.18 |

